# Cortico-striatal beta-oscillations as a marker of learned reward value

**DOI:** 10.1101/2022.10.24.513264

**Authors:** M.F. Koloski, S. Hulyalkar, T. Tang, X. Wu, L. Fakhraei, S.A. Barnes, J. Mishra, D.S. Ramanathan

## Abstract

Single neuron correlates of reward value have been observed in brain regions along the cortico-striatal pathway including ventral striatum, orbital, and medial prefrontal cortex. Brain imaging studies in humans further validate these findings and suggest that value is represented in a network of brain regions opposed to a particular area. Neural activity oscillates at periodic frequencies to coordinate long-range communication in widespread, dynamic networks. To explore how oscillatory dynamics across brain regions may represent reward value, we measured local field potentials of male Long-Evans rats during three distinct behavioral tasks, each probing a different aspect of reward processing. Our goal was to use a data-driven approach to identify a common electrophysiology property associated with reward value. We found that reward-locked oscillations at beta frequencies, in both single units and local field potentials, were markers of positive reward valence. More importantly, Reward-locked beta-oscillations scaled with expected reward value on specific trial types and in a behaviorally relevant way across tasks. Oscillatory signatures of reward processing were observed throughout the cortico-striatal network including electrodes placed in orbitofrontal cortex, anterior insula, medial prefrontal cortex, ventral striatum, and amygdala. These data suggests that beta-oscillations reflect learned reward value in a distributed network, and this may serve as a stable and robust bio-marker for future studies.

## Introduction

Reward processing comprises the set of neural systems associated with appetitive, motivational, or pleasurable stimuli (1, 2). Deficits in reward-processes are linked with learning and decision-making impairments and likely contribute to anhedonia, amotivation, and substance abuse problems observed in various psychiatric conditions (1, 3). Thus, identifying preclinical bio-makers of reward processing will help assess behavioral deficits and expand treatment options that are currently limited (2–5).

Past studies highlight the relevance of cortico-striatal circuitry for reward learning. The ventral striatum, and in particular the nucleus accumbens, is connected to the medial prefrontal cortex, orbitofrontal cortex, and basolateral amygdala through cortico-striatal-limbic reward-network projections (1,2,6–14). This extended “reward” network is innervated by midbrain dopamine neurons originating from the ventral tegmental area, which contribute to reward processing behaviors through reward-prediction error signals (the difference between expected and actual rewards) (10,12,15–19). Thus, standard models of reinforcement/reward learning posit that dopamine neurons carry a “RPE” signal that then modulates distinct parts of the cortico-striatal reward network in specific ways.

Single-unit activity is high-dimensional. Neurons from any brain region can encode a diverse array of task-related processes (20–23). For example, single neurons in ventral striatum, prefrontal and orbitofrontal cortex can be modulated both during reward anticipation and delivery (13,23–29), and can be modulated by different types (6, 30), magnitudes (6,24,31,32), and locations of reward (28). Low-dimensional representations of population activity provide a more robust, stable, and simpler framework to identify neuro-behavioral relationships and can be compared with human neuroimaging data (20,21,33). Local field potentials (LFP) offer an opportunity to bridge micro- and macroscopic levels of brain activity and (in the correct circumstance) can reflect low-dimensional population level features of single-units(22,33–37).

We have previously used multi-site LFP recordings to characterize networks operating at distinct oscillatory frequencies to support behavioral inhibition and default-mode-like processing (38, 39). Here we utilize our multi-site LFP approach to identify electrophysiology markers linked with reward expectation and outcome. It is unclear whether a neural signature may be unique to a specific domain of reward processing or may represent a common substrate across domains. Therefore, to increase the behavioral specificity of our electrophysiological markers, we examined data from three distinct behavioral tasks (with different animals trained up on each task). The behavioral tasks used each contribute a separate dimension of reward learning (**Table 1**): 1) A go/wait behavioral inhibition task was used to identify signals related to the valence of feedback (reward vs. no reward), with reward essentially scaled to performance; 2) A temporal discounting task was used to identify how reward-locked signals scale with subjective value of both reward magnitude (high vs. low reward) and temporal delay (0.5 to 20s); 3) Finally, a probabilistic reversal learning task was used to identify signals modulated by the learned probability of reward delivery (high vs. low-probability).

**Table 1:** The experimental design including number of subjects, behavioral session, time windows of interest and contrasts for analysis are provided for each of the three behavioral tasks.

|  | **Go/Wait Behavioral Inhibition** | **Temporal Discounting** | **Probabilistic Reversal Learning** |
| --- | --- | --- | --- |
| Local Field Potential Recordings |  |  |  |
| Subjects | 12 | 10 | 7 |
| with histology | 6 | 9 | 7 |
| Behavioral Sessions | 67 | 124 | 79 |
| with electrophysiology | 67 | 124 | 36 |
| Time Window | 500 – 2500ms after response | 0-1000ms after reward onset | 500- 2500ms after response |
| Contrasts | go-cue vs. wait-cue trials<br>reward vs. no reward | high vs. low reward magnitude<br>temporal delay<br>(0.5, 1, 2, 5, 10, 20s) | high vs. low probability of reward<br>reward vs. no reward |
| Single Unit Recordings |  |  |  |
| Subjects | 8 |  |  |
| with histology | 5 |  |  |
| Behavioral Sessions | 62 |  |  |
| with electrophysiology | 62 |  |  |
| Time Window | 0 – 2000ms after response |  |  |
| Contrasts | Go-Cue vs. Wait-Cue Trials<br>Reward vs. No Reward |  |  |

On each task, we first examined activity in the lateral orbitofrontal cortex (lOFC), a cardinal brain region consistently identified for its role in evaluating reward outcomes and expectancies to drive adaptive behavior (1,7,26,40–42). Next, we analyzed pertinent oscillatory markers from 12 of our 32 electrodes that were placed in areas along the cortico-striatal pathway in brain regions previously identified as inter-connected with the ventral striatum. We provide evidence of oscillatory activity at beta and high-gamma frequencies found consistently across our three tasks that modulates with expected reward value.

## Results

### Beta Frequency Oscillations Linked with Positive Valence Feedback

A full description of behavior on the go/wait task in animals with LFP probes can be seen in our prior publication describing inhibition and stimulus-response oscillatory signatures (39). Animals were shown two visual stimuli, one which required an immediate response (go-cued trials) and the other which required the animal to withhold from responding for 2s (wait-cued trials) (**Fig. 1A**). Across behavioral sessions, animals performed better on go-cued trials compared to wait-cued trials (**Fig. 1B**). Animals could generally distinguish between go and wait-cues indicated by a significant difference in reaction times (*t*_(61)_ =17, *p<.001*) and greater accuracy on go-cued trials (*t*_(61)_ =18, *p<.001*). On go-cued trials animals had a mean reaction time of 610 +/− 160ms and correctly responded within 2s of the visual cue on 94.0 +/− 9.2% of trials (data averaged across 62 sessions from 12 animals). On wait-cued trials animals took longer to respond (1700 +/− 460ms) and were able to correctly wait on 41.0 +/− 23.0% of trials **(Fig. 1B)**.

**Figure 1:**
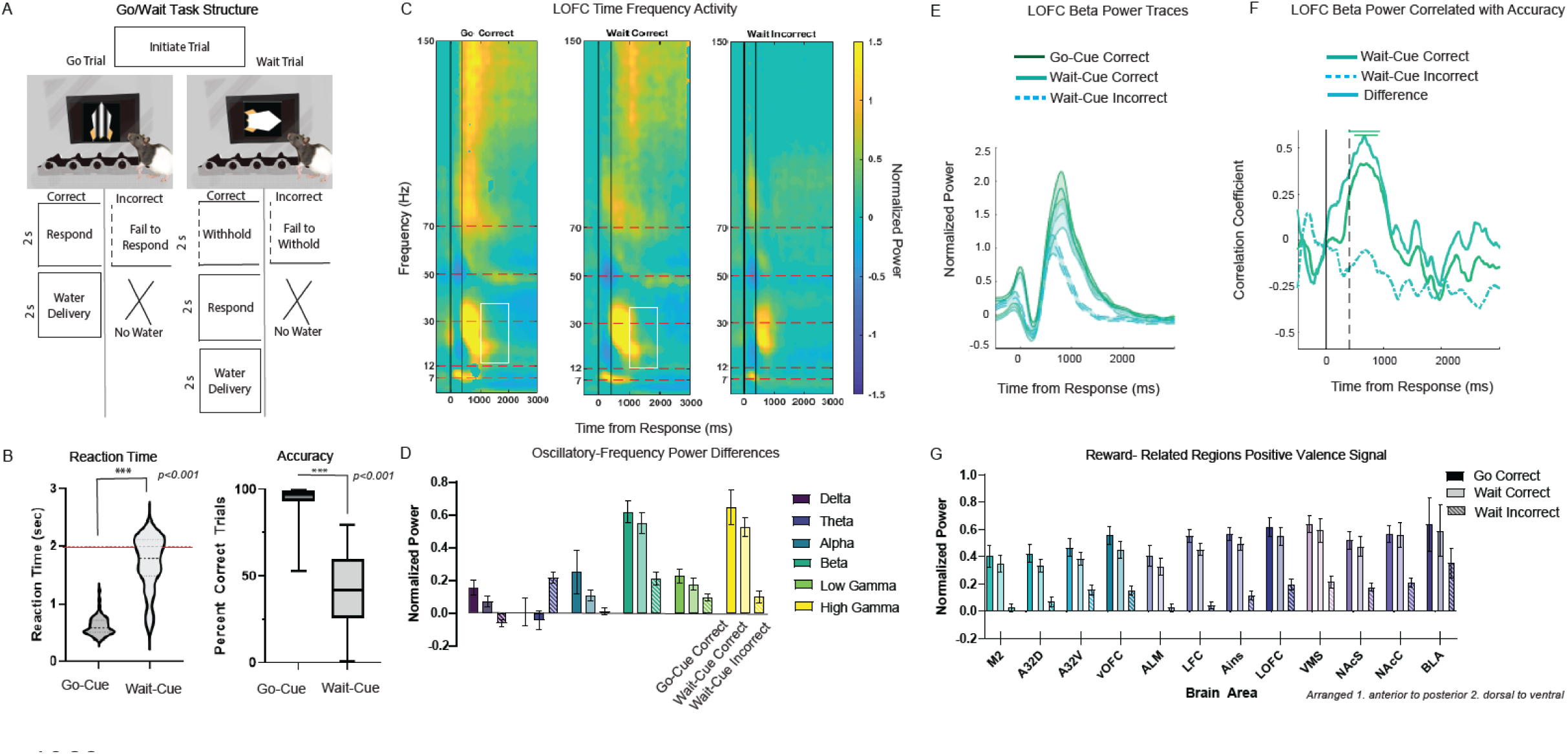
Positive Valence Representation on the Response Inhibition Task. (A) Trial structure of the go/wait task. Animals were shown two visual stimuli: a striped vertical rocket indicated a go-cue trial (respond within 2s) and a solid horizontal rocket denoted a wait-cue (withhold from responding for 2s). On correct trials rats were given 2s (10µL/s) of water. (B) Behavior on the go/wait task of animals with LFP implants (N= 67 sessions). Violin plot shows the median (thick dotted line), interquartile range (thin dotted line), and distribution shape of reaction times (s) on go-cue and wait-cue trials. Red horizontal line represents the 2s time to response on go-cue trials and time needed to withhold on wait-cue trials. Bar plots shows the mean and SEM of proportion correct trials on go-cue (respond within 2s) and wait-cue (withhold for 2s) trials. Dots show individual values per session. (C) Time-frequency plots of z-scored lOFC power time-locked to response for frequencies ranging from 0-150 Hz on correct go-cue correct (reward), wait-cue correct (reward) and wait-cue incorrect (no reward) trials. Vertical lines represent the response time and reward onset time. (D) Average lOFC power across delta (1-4 Hz), theta (4-8 Hz), alpha (8-12 Hz), beta (15-30 Hz), and low gamma (50-70 Hz) and high gamma (70-150 Hz) frequencies during the reward-feedback period (500-2500ms after response). We used a linear mixed model to quantify differences in power across frequency bands on the LOFC electrode for go-cue correct (dark), wait-cue correct (light) and wait-cue incorrect trials (striped). Mean and SEM are plotted. (E) Line plots show mean (middle line) and SEM (outer boundaries and shaded region) beta power on lOFC electrode time-locked to response onset for go-cue correct (dark), wait-cue correct (light), and wait-cue incorrect trials (dashed). (F) Line plots show correlation between lOFC beta power and wait trial accuracy. Separate lines are plotted for wait-cue correct (dark), wait-cue incorrect (dashed), and the difference between correct and incorrect trials (light). (G) Beta power from 12 putative reward-related brain regions on go-cue correct (dark), wait-cue correct (light) and wait-cue incorrect trials (striped) during the reward-feedback period (1550ms-2550ms). The mean and SEM for each trial type and brain region of interest are shown. Brain areas are shown in order from 1. anterior to posterior and 2.dorsal to ventral.

For all tasks, we began by analyzing electrophysiological activity, time-locked to the response, from lateral OFC (lOFC)-a cardinal reward processing brain region in the cortico-striatal network. Then, after identifying pertinent oscillatory frequency bands of interest during reward-feedback, we performed a second linear mixed model to analyze 12 electrodes in the cortico-striatal network (**Table 2**). The primary goal of our first analysis was to identify electrophysiological markers that differentiated between positive and negative feedback on the go/wait task. Feedback on correct trials consisted of water delivery 400ms after the response at the rate of 10µl/sec, for a duration of 2s. Feedback on incorrect trials consisted of a 5s flashing house-light and an auditory 1000Hz tone with no water delivery. We analyzed mean time-frequency (TF) power across sessions (N=62) from correct go-cued trials (animals received a go-cue and responded within two seconds), correct wait-cued trials (animals received a wait-cue and waited two seconds before responding) and incorrect wait-cued trials (animals received a wait-cue but failed to wait two second before responding). Due to the high accuracy on go-cued trials (**Fig. 1B**), there were very few incorrect go-cued trials (failing to respond within two seconds) and thus we did not analyze this trial type in the subsequent analyses. In the first linear mixed model, we took the average power across delta (1-4 Hz), theta (4-8 Hz), alpha (8-12 Hz), beta (15-30 Hz), low gamma (50-70 Hz) and high gamma (70-150 Hz) frequencies during a two second reward-feedback window from 500-2500ms after response (corresponding to the timepoint of reward delivery) on the lOFC electrode (**Table 3**).

**Table 2:** Electrode sites of interest are listed in order from 1. Anterior to possterior 2. Dorsal to ventral.

| Abbreviation | Brain Area |
| --- | --- |
| M2 | Secondary Motor Cortex |
| A32D | Dorsomedial Prefrontal Cortex |
| A32V | Ventromedial Prefrontal Cortex |
| vOFC | Ventral Orbitofrontal Cortex |
| ALM | Anterolateral Motor Cortex |
| LFC | Lateral Frontal Cortex |
| Ains | Anterior Insula |
| lOFC | Lateral Orbitofrontal Cortex |
| VMS | Ventromedial Striatum |
| NAcS | Nucleus Accumbens Shell |
| NAcC | Nucleus Accumbens Core |
| BLA | Basolateral Amygdala |

**Table 3:**
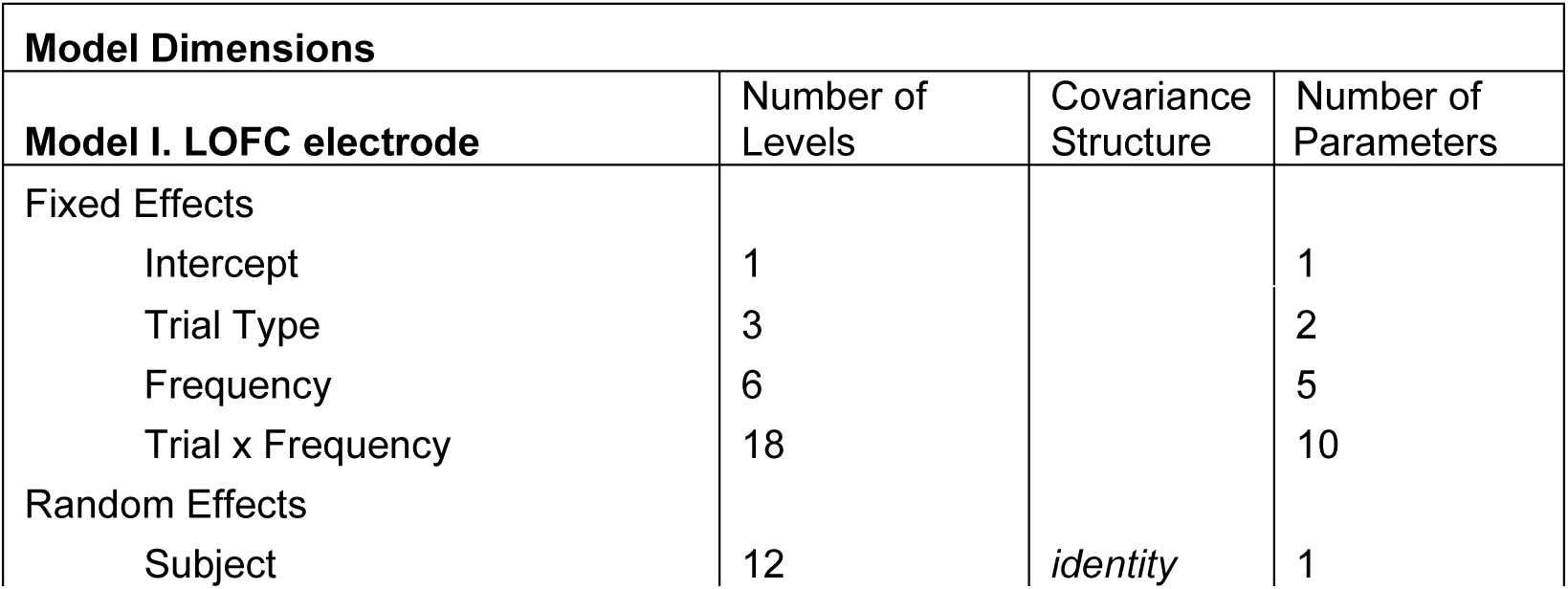
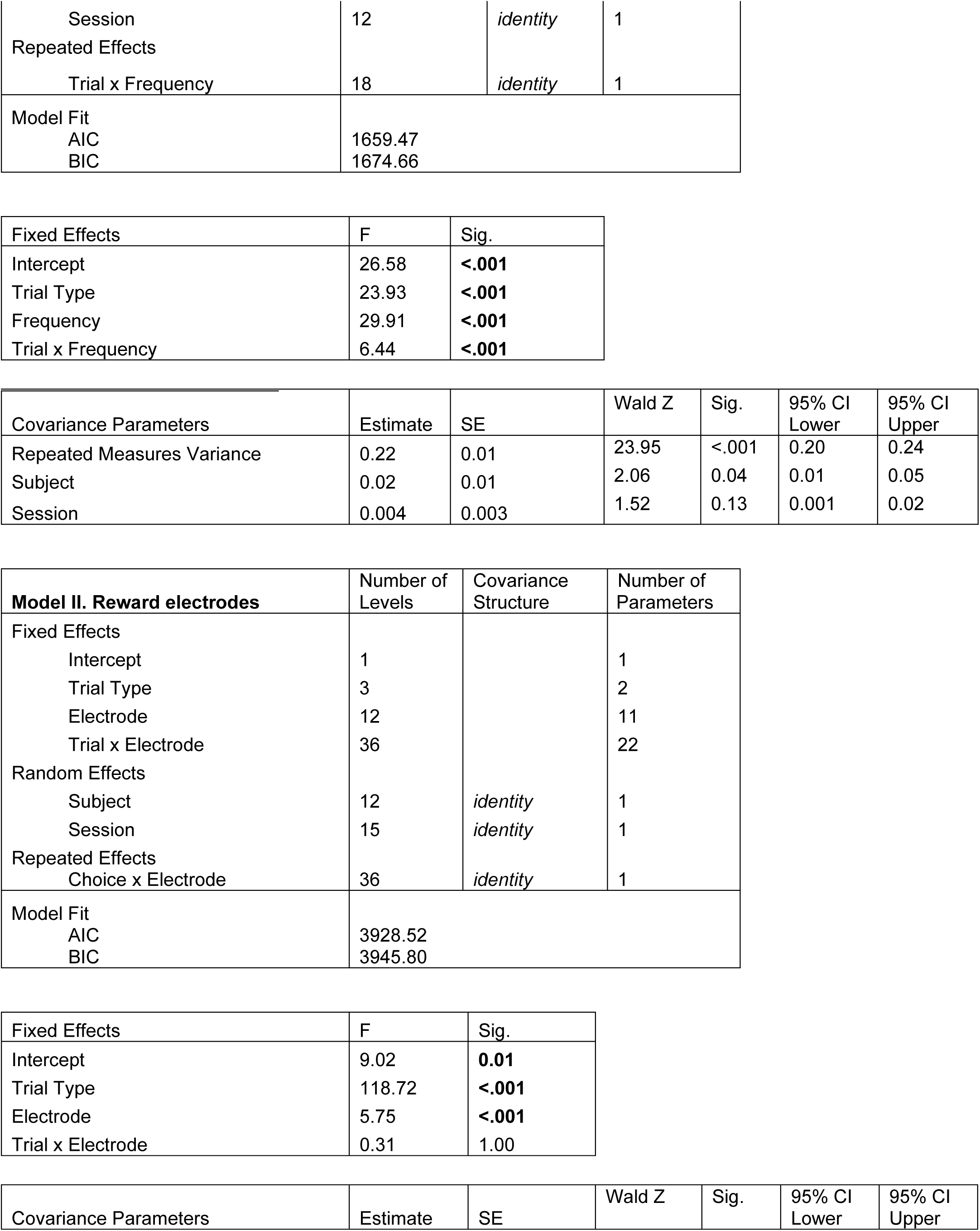
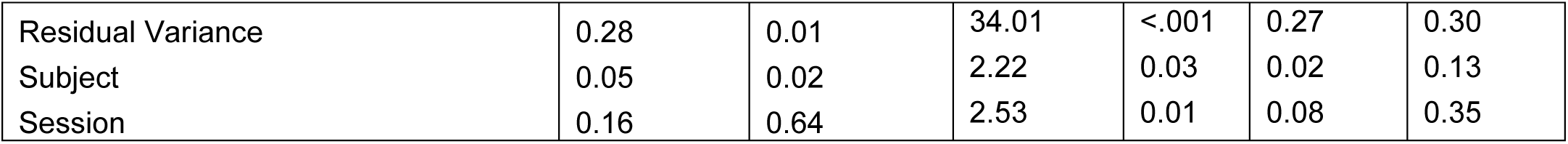
Linear mixed model design, fixed effects, and covariance parameter to explore power differences during reward outcome on the go/wait inhibition task.

| Model Dimensions |  |  |  |
| --- | --- | --- | --- |
| Model I. LOFC electrode | Number of Levels | Covariance Structure | Number of Parameters |
| Fixed Effects |  |  |  |
| Intercept | 1 |  | 1 |
| Trial Type | 3 |  | 2 |
| Frequency | 6 |  | 5 |
| Trial x Frequency | 18 |  | 10 |
| Random Effects |  |  |  |
| Subject | 12 | *identity* | 1 |
| Session | 12 | *identity* | 1 |
| Repeated Effects |  |  |  |
| Trial x Frequency | 18 | *identity* | 1 |
| Model Fit |  |  |  |
| AIC | 1659.47 |  |  |
| BIC | 1674.66 |  |  |

|  |  |  |
| --- | --- | --- |
| Fixed Effects | F | Sig. |
| Intercept | 26.58 | **<.001** |
| Trial Type | 23.93 | **<.001** |
| Frequency | 29.91 | **<.001** |
| Trial x Frequency | 6.44 | **<.001** |

|  |  |  |  |  |  |  |
| --- | --- | --- | --- | --- | --- | --- |
| Covariance Parameters | Estimate | SE | Wald Z | Sig. | 95% CI Lower | 95% CI Upper |
| Repeated Measures Variance | 0.22 | 0.01 | 23.95 | <.001 | 0.20 | 0.24 |
| Subject | 0.02 | 0.01 | 2.06 | 0.04 | 0.01 | 0.05 |
| Session | 0.004 | 0.003 | 1.52 | 0.13 | 0.001 | 0.02 |
| **Model II. Reward electrodes** | Number of Levels | Covariance Structure | Number of Parameters |
| Fixed Effects |  |  |  |
| Intercept | 1 |  | 1 |
| Trial Type | 3 |  | 2 |
| Electrode | 12 |  | 11 |
| Trial x Electrode | 36 |  | 22 |
| Random Effects |  |  |  |
| Subject | 12 | *identity* | 1 |
| Session | 15 | *identity* | 1 |
| Repeated Effects |  |  |  |
| Choice x Electrode | 36 | *identity* | 1 |
| Model Fit |  |  |  |
| AIC | 3928.52 |  |  |
| BIC | 3945.80 |  |  |

|  |  |  |
| --- | --- | --- |
| Fixed Effects | F | Sig. |
| Intercept | 9.02 | **0.01** |
| Trial Type | 118.72 | **<.001** |
| Electrode | 5.75 | **<.001** |
| Trial x Electrode | 0.31 | 1.00 |

|  |  |  |  |  |  |  |
| --- | --- | --- | --- | --- | --- | --- |
| Covariance Parameters | Estimate | SE | Wald Z | Sig. | 95% CI Lower | 95% CI Upper |
| Residual Variance | 0.28 | 0.01 | 34.01 | <.001 | 0.27 | 0.30 |
| Subject | 0.05 | 0.02 | 2.22 | 0.03 | 0.02 | 0.13 |
| Session | 0.16 | 0.64 | 2.53 | 0.01 | 0.08 | 0.35 |

We examined fixed effects of trial type (go-correct, wait-correct, wait-incorrect), frequency, and their interaction. We examined random effects of subject and session. We found a main effect of trial type (F_(2,1147.03)_=23.93, *p<.001*), main effect of frequency (F_(5,1147.03)_=29.91, *p<.001*), and a significant interaction between frequency and trial type (F_(10, 1147.03)_ =6.44, *p<.001*). Post-hoc analyses (Bonferonni corrected) revealed that the main effect of frequency was driven by greater power on the lOFC electrode during reward-feedback at beta (EMM= 0.50, SEM= 0.06, CI= 0.38, 0.61) and high-gamma frequencies (EMM= 0.46, SEM= 0.06, CI= 0.34 0.58). (**Fig. 1C;D**). Importantly, oscillatory activity at beta and high-gamma frequencies was different based on trial type. Beta power was greater on rewarded trials (go-cue correct trials: EMM= 0.66, SEM= 0.07, CI= 0.12, 0.41; wait-cue correct trials: EMM= 0.59, SEM= 0.07, CI= 0.44, 0.73), compared to unrewarded/ incorrect wait-cue trials (EMM= 0.24, SEM= 0.07, CI= 0.1, 0.39) (**Fig. 1C;D**). On rewarded trials the peak beta activity in lOFC occurred at 805ms after the response (~400ms after reward onset) and lasted for around 2 seconds-the approximate time of reward delivery (**Fig. 1E**). In this mixed effects model, the two random effects were subject and session. Subject contributed to 8.2% of variance and was significant according to a WaldZ metric (Wald Z= 2.06, *p=0.04*) (**Supp Fig. 1**). Session accounted for only 1.7% of variance in the model and was not a significant contributor.

Dopaminergic signals related to reward are often linked with a “reward-prediction-error”, i.e. they are typically positively modulated by difference between expectation of reward and reward delivery. By contrast, single neurons within OFC has been observed to predict the opposite of an RPE – i.e. they are related to reward-prediction (43). To understand whether lOFC beta-power was linked with reward-prediction vs. an RPE on this task, we focused on whether the average beta-power during the wait-cue trials was linked with accuracy on that session using a linear regression analysis between session performance and mean session beta-power. We hypothesized that, if related to an RPE, beta power would be negatively correlated with wait-cue accuracy whereas if it was related to reward prediction it should be positively correlated with performance. We found that lOFC beta power was significantly positively correlated with wait accuracy (FDR corrected) from 500-1000ms on wait-cue rewarded trials. The difference between the correct and incorrect trials for the wait-cue also predicted greater accuracy on wait-cued trials. (**Fig. 1F**). This relationship importantly indicated two things: 1) beta-power is unlikely to be a trivial artifact or related to noise, as noise wouldn’t obviously be correlated with performance; 2) lOFC beta-power was a marker of reward-prediction and not RPE.

To better understand the spatial distribution of the reward-prediction beta activity beyond lOFC, we next analyzed beta power from 12 of our 32 electrodes (M2, A32D, A32V, vOFC, ALM, LFC, Ains, lOFC, VMS, NAcS, NAcC, BLA) (**Table 2**), chosen based on cortico-striatal regions connected with ventral striatum for whom we had electrode locations and were large enough regions to make LFP a meaningful measure. We examined LFP activity across divisions of medial prefrontal cortex, orbitofrontal cortex, ventral striatum, anterior insula, and basolateral amygdala. As seen on the lOFC electrode, there was a main effect of trial type (F_(2,2313.76)_=118.72, *p<0.001*) on beta frequency power during reward-feedback (**Fig. 1G**). There was also a main effect of electrode (F_(11,2313.76)_=5.75 *p<0.001*) but no significant trial x electrode interaction (F_(22,231.76)_=0.31, *p=1.0*). Post-hoc (Bonferroni corrected) tests revealed the main effect of electrode was driven by increased power on the BLA electrode (EMM= 0.53, SEM= 0.13, CI= 0.26, 0.79) that was greater for rewarded vs. non-rewarded trial types. LOFC (EMM= 0.45, SEM-0.13, CI= 0.19, 0.72) and VMS (EMM= 0.48, SEM= 0.13, CI= 0.21, 0.75) also had increased beta-frequency power on rewarded trials (**Fig. 1G**). Subjects contributed to 11% of variance in the model, which was significant according to a Wald Z metric (Wald z = 2.22, *p=0.03*). Session contributed to 32.6% of variance in the model which was also significant (Wald z= 2.526, *p=0.01*).

### Beta-Oscillations Related to Single-unit Activity in OFC During Reward Feedback

Despite the name, “local” field potentials are challenging to properly localize (35,44,45). To better understand whether the beta frequency activity observed during positive reward-feedback was related to local spiking activity within a particular brain region, we recorded single-units from the OFC of 8 different male Long-Evans rats performing the go/wait task (**Table 1**). The version of the task used for single-unit recordings was slightly modified (due to a slight change in coding up of this version of the task), from a 400ms delay between response and reward as noted above, to only a 30ms delay. As we were focused on the period post-feedback and not during the anticipation period, this modification did not affect task performance. We recorded 376 neurons across 62 sessions (5.81 units +/− 0.03 per session). While performance on the task was worse in these animals compared to those with LFP implants, rats were still able to discriminate between go-cue and wait-cue trials. Reaction time was different between trial types (*t*_(61)_=7.8, *p<.001*): 800 +/− 180ms on go-cue trials and 1100 +/− 330ms on wait-cue trials. Accuracy on go-cue trials was 73.0 +/−24.0%, compared to 30.0 +/− 20.0% on wait-cue trials (*t*_(61)_=8.7, *p<.001*) (**Fig. 2A**).

**Figure 2:**
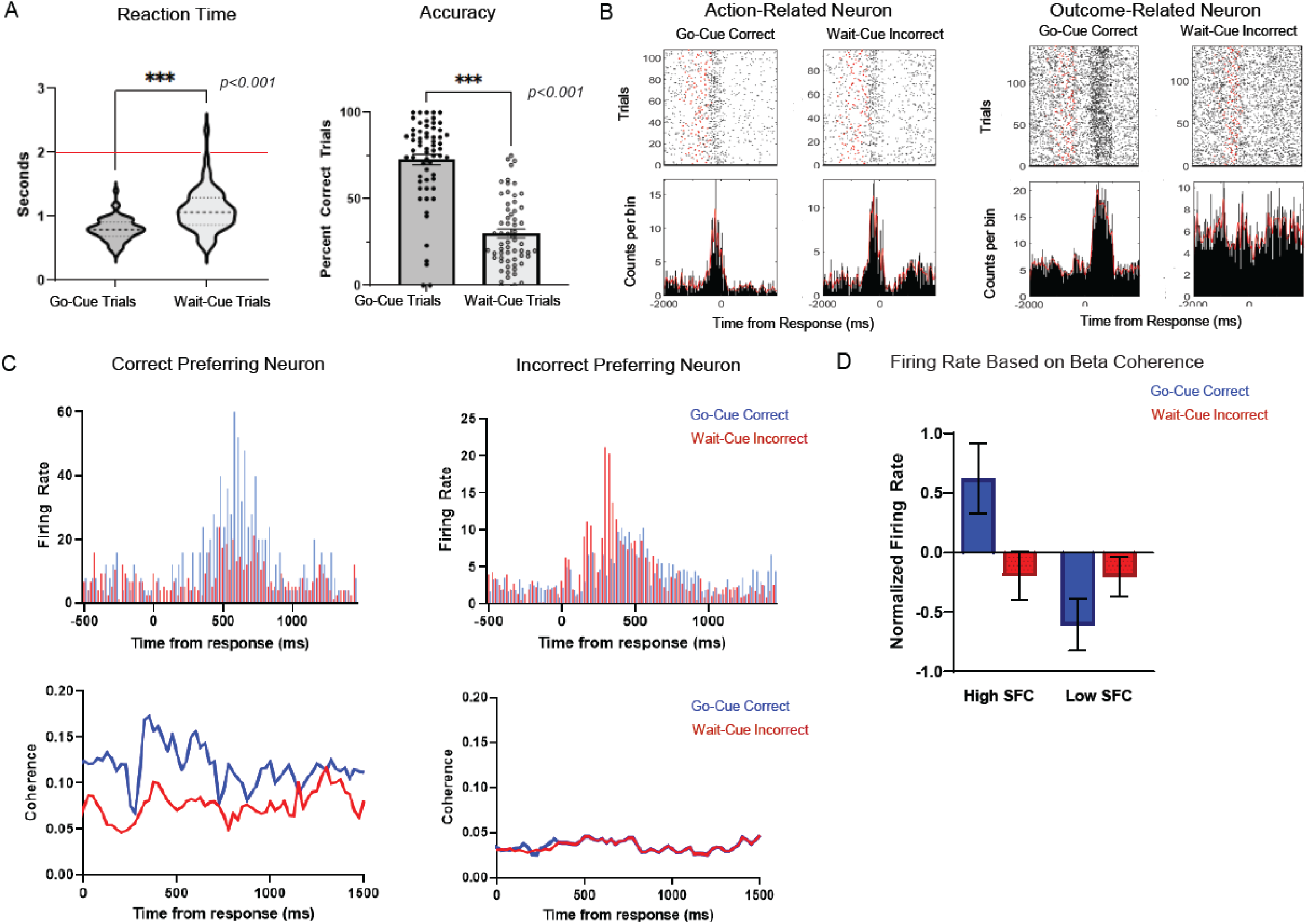
Single-Units Related to Positive Valence Reward Outcome. (A) Behavior on the go/wait task of animals with single unit implants (N= 62 sessions). Violin plot shows the median (dark dotted line), interquartile range (light dotted line), and distribution shape of reaction times (s) on go-cue trials and wait-cue trials. Red line drawn at 2s represents the time to respond on go-cue trials and the time needed to withhold on wait-cue trials. Bar plots show proportion of correct trials on go-cue and wait-cue trials. Dots show individual values per session. (B) Individual examples of an action-related neuron (peak firing rate increases or decreases prior to the response) and an outcome-related neuron (peak firing rate increases or decreases after the response). Activity of each neuron is plotted for go-cue correct (reward) and wait-cue incorrect (no reward) trials time-locked to the response (time 0). Raster plots (top-panel) show spiking activity across trials (each horizontal line). Red dots indicate trial onset. Peri-event stimulus histograms (bottom panel) show firing rate (counts per bin) time-locked to response. Red lines indicate the mean activity of the unit across trials. (C) Individual examples of a correct preferring (more activity on rewarded trials) and an incorrect preferring (more activity on non-rewarded trials) unit. Firing rate (counts/bin) from go-cue correct (blue) and wait-cue incorrect (red) trials are plotted on top of each other time-locked to response for comparison (top-panel). Examples of beta-frequency SFC are shown for the same correct and incorrect preferring neurons on go-cue correct (blue) and wait-cue incorrect (red) trials (bottom-panel). Coherence values are time-locked to response. (D) Normalized firing rate (spikes/s) of neurons split into “high” or “low” groups based on their beta-SFC values during reward-feedback. Firing rates are plotted separately for go-cue correct (blue) and wait-cue incorrect (red) trials. Error bars indicate SEM.

After excluding sessions with a limited trial number (< 30 trials) and units with low firing rates (< 2 spikes/s), 228 units were included for subsequent analyses. 125 neurons (33%) were defined as task-modulated based on our criteria of an increase/decrease of two standard deviations above baseline for >75 consecutive ms. This included single units with both peak firing rate increases or decreases that occurred both prior to the response (action-related) or after the response (outcome or feedback-related) (**Fig. 2B**). The average peak firing rate activity of action-related neurons was 375ms before the response (time 0). The average peak firing rate of outcome-related neurons was 225ms after response (~195ms after reward onset). 103 neurons were not task-modulated based on our criteria.

Our main goal for studying single units was to determine whether they were modulated by reward-related beta-oscillations. We first used spike-field-coherence (SFC) to assess the relationship between spiking and oscillatory activity during the reward-feedback period (0 to 2000ms after response) (44,46–48). Units with significant task-related suppression or missing LFP data stream were not included in SFC analysis (173 units remaining). We observed that neurons with greater firing rate on rewarded, go-cue correct trials compared to non-rewarded, wait-cue incorrect trials (“correct preferring”) showed increased beta frequency SFC modulation on correct trials vs. incorrect during reward-feedback (example neuron, **Fig. 2C**). By contrast, outcome-related neurons with greater firing rate on wait-cue incorrect trials (“incorrect preferring”) did not show as great of SFC modulation at beta frequencies (example neuron, **Fig. 2C**).

We grouped neurons into two categories solely based on their beta SFC value during the reward-feedback period, and measured how this grouping was linked with firing rate for both correct and incorrect trials. The “high-SFC” neurons were identified as neurons with one standard deviation higher-than-average beta-SFC; and “low-SFC” neurons were identified as neurons with one standard deviation lower-than-average beta-SFC. We found a main effect of SFC category (high vs. low) (F_(1,314)_ 5.11, p*=0.024*) on reward-feedback firing rate, and a significant interaction (F _(1,314)_ 4.45, p*=0.036*) between SFC category and trial type (go-cue correct vs. wait-cue incorrect) (**Fig. 2D**). Neurons in the “high” SFC group had an average firing rate of 0.66 +/−0.41 spikes/s on go-cue correct trials compared to the “low” SFC group neurons which had an average firing rate of −0.78+/−0.30 spikes/ s. On wait-cue incorrect trials, firing rate was not modulated based on SFC value. Neurons in the “high” SFC group had an average firing rate of −0.20 +/− 0.27 spikes/s on wait-cue incorrect trials and “low” SFC neurons had an average of −0.15 +/− 0.17 spikes/s. The firing rate of “high” and “low” SFC neurons was similar on non-rewarded (wait-cue incorrect) trials but was significantly different on rewarded (go-cue correct) trials (mean difference [high-low]= 1.44, *corrected p=0.004*) (**Fig. 2D**). Thus, we found that single-units from OFC with higher reward-locked beta SFC are also more likely to be positively modulated by reward while those with low reward-locked SFC are more likely to be suppressed by rewards.

### Beta Power Reflects Dimensions of Reward Prediction and Value

Our data from the go/wait task suggested that beta activity within lOFC and other cortico-striatal regions relates to positive valence (i.e. rewards) and may relate to reward prediction or expected value. However, as this task was not specifically designed to modulate aspects of reward value, it was still possible that, on a different task, we would see a different relationship between beta oscillations and reward. Using a new group of animals to study reward prediction signals on a different task allows for replication and to rule out beta as related to some non-specific aspects of reward consumption unrelated to subjective value or prediction. To further explore these hypotheses, we recorded LFP activity on a new set of animals (N=10) trained to perform a temporal discounting task (**Fig. 3A**). On this task, animals were given the choice of a low-value reward delivered with a fixed delay of 500ms after response or a higher-value reward delivered at variable delays of between 500ms to 20 seconds. To allow for greater numbers of trials for electrophysiological analysis, delays on the high-reward condition were kept constant throughout each session but varied across sessions. Low-value rewards consisted of 10ul whereas high-value rewards were 30ul (both delivered at a rate of 10 ul/sec). Previous work suggests reward value is negatively influenced by temporal costs associated with earning a reward (19,31,49–51). In the context of this task, the subjective value of the high reward choice decreases as the length of the delay required to obtain reward (temporal cost) increases. Results from the temporal discounting task are based on 124 total sessions (average for each rat was 2 sessions/ variable delay) (**Table 1**). As expected, animals’ preference shifted from high-value choice to the low-value choice as the delay to reward increased (F_(5,45)_ =30.9, *p =<0.001,* two-way ANOVA) (**Fig. 3B**). When delays of each choice were the same (500ms), animals strongly prefer the high-value (30ul) reward (90.4 +/− 1.4 % high-value choices per session). When the high-value reward follows a 20s delay, rats only select the high reward choice 24.2 +/− 6.8% of trials, showing a clear preference for the immediate, low-value reward. We do see individual differences emerge in the average rate of discounting across delays (F_(9,45)_=7.02, *p<0.001,* two-way ANOVA) (**Fig. 3B**).

**Figure 3:**
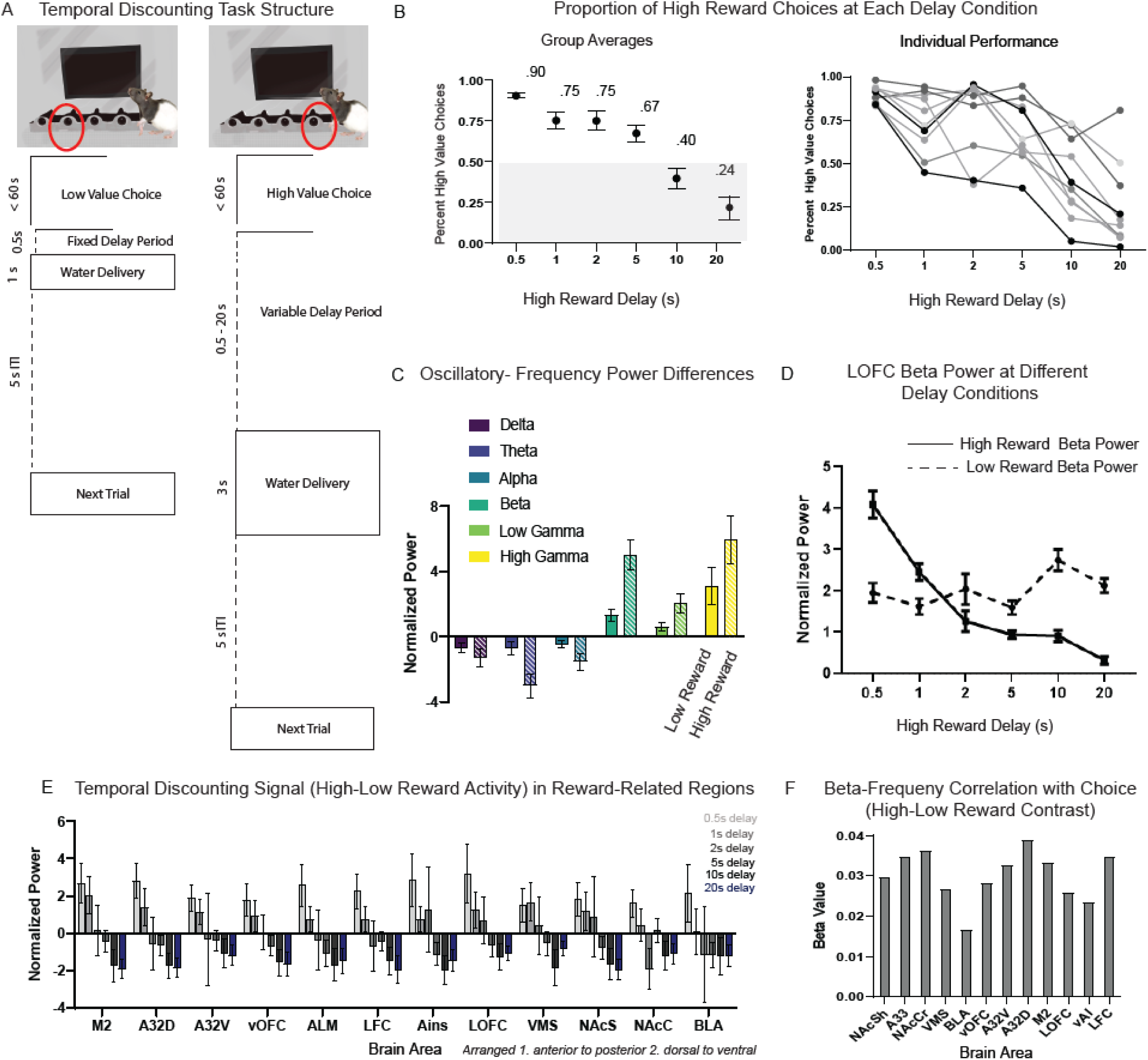
Subjective Value Representation on the Temporal Discounting Task. A) Trial structure of the temporal discounting task. Animals were given the choice of a low magnitude reward (10µL) delivered immediately (delay of 500ms after response), or a high magnitude reward (30µL) delivered at variable delays of 0.5s up to 20s. Delays on the high-value choice were kept constant throughout each session but varied across sessions. (B) Behavior on the temporal discounting task shown as proportion of high-value choices per session (N=124 sessions). The group mean and SEM are shown at each delay condition (0.5, 1, 2, 5, 10, and 20s). A horizontal line is drawn at 0.5 to indicate when proportion of high-value choices fall below the 50/50 mark. The average proportion of high-value choices at each delay condition is also plotted separately for each rat to show individual differences in discounting rates. (C) Average power across delta (1-4 Hz), theta (4-8 Hz), alpha (8-12 Hz), beta (15-30 Hz), low gamma (50-70 Hz) and high gamma (70-150Hz) frequencies on the lOFC electrode one second after reward onset on equal low value and high value delay sessions (0.5s). Error bars represent SEM. (D) The normalized beta power on the lOFC electrode at each high-value delay condition (0.5, 1, 2, 5, 10, 20s). Power is averaged across the first second following reward delivery and the mean/SEM are plotted separately for high-reward choices (solid line) and low-reward choices (dashed line). (E) The difference in power on high reward and low reward choices (high-value choice – low-value choice) across 12 brain regions. Beta power during reward-feedback (averaged activity one second after response) is plotted at each delay condition (0.5, 1, 2, 5, 10, 20s) for each brain region. Brain regions are organized from 1. anterior to posterior and 2. dorsal to ventral. (G) Beta values from the logistic regression analysis for 12 brain regions using power for the difference of trial types (high-value – low-value). Asterisks indicate brain regions with significant p-values after FDR-correction.

The first question we asked was whether lOFC beta power is modulated by expected reward value. Specifically, we hypothesized that if beta reflects an aspect of expected reward value, then power should be greater for the high (30ul) compared to the low (10ul) reward magnitude when delays were the same (500ms for both). We analyzed only the first second of activity post-reward to ensure that, for both trial types, animals were receiving the same quantity of reward during the period of analysis (i.e., during the first second of reward deliver for both reward types there was an equivalent reward delivery). Using a linear mixed model to account for subject and session variance, we first investigated data across all frequencies (delta: 1-4 Hz; theta: 4-8 Hz; alpha: 8-12 Hz; beta: 15-30 Hz; low gamma: 50-70 Hz; and high gamma: 70-150 Hz) at the lOFC electrode. Our model measured base-line normalized power modulation (the ratio of activity at a particular time point relative to base-line) as the dependent variable across different frequency bands (delta, theta, alpha, beta, low gamma, high gamma) and trial type (high or low reward choice) with subject and session variance as random effects (**Table 4**).

**Table 4:**
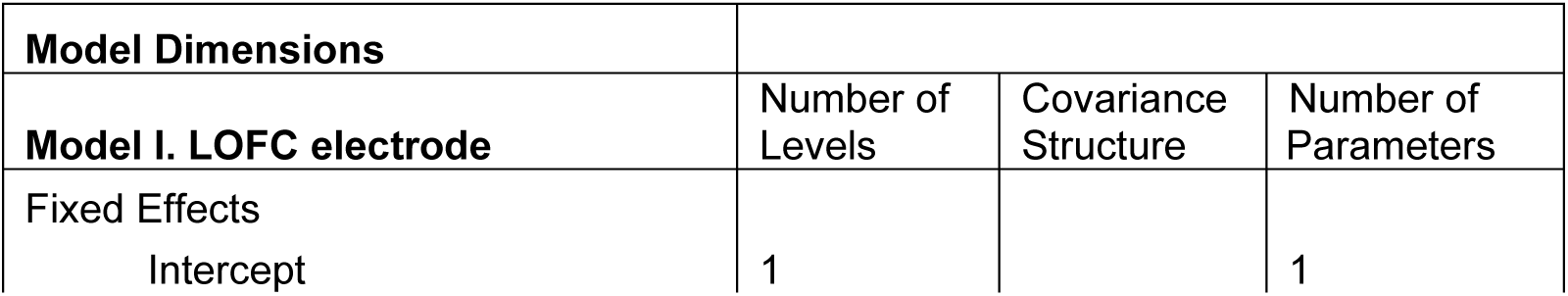
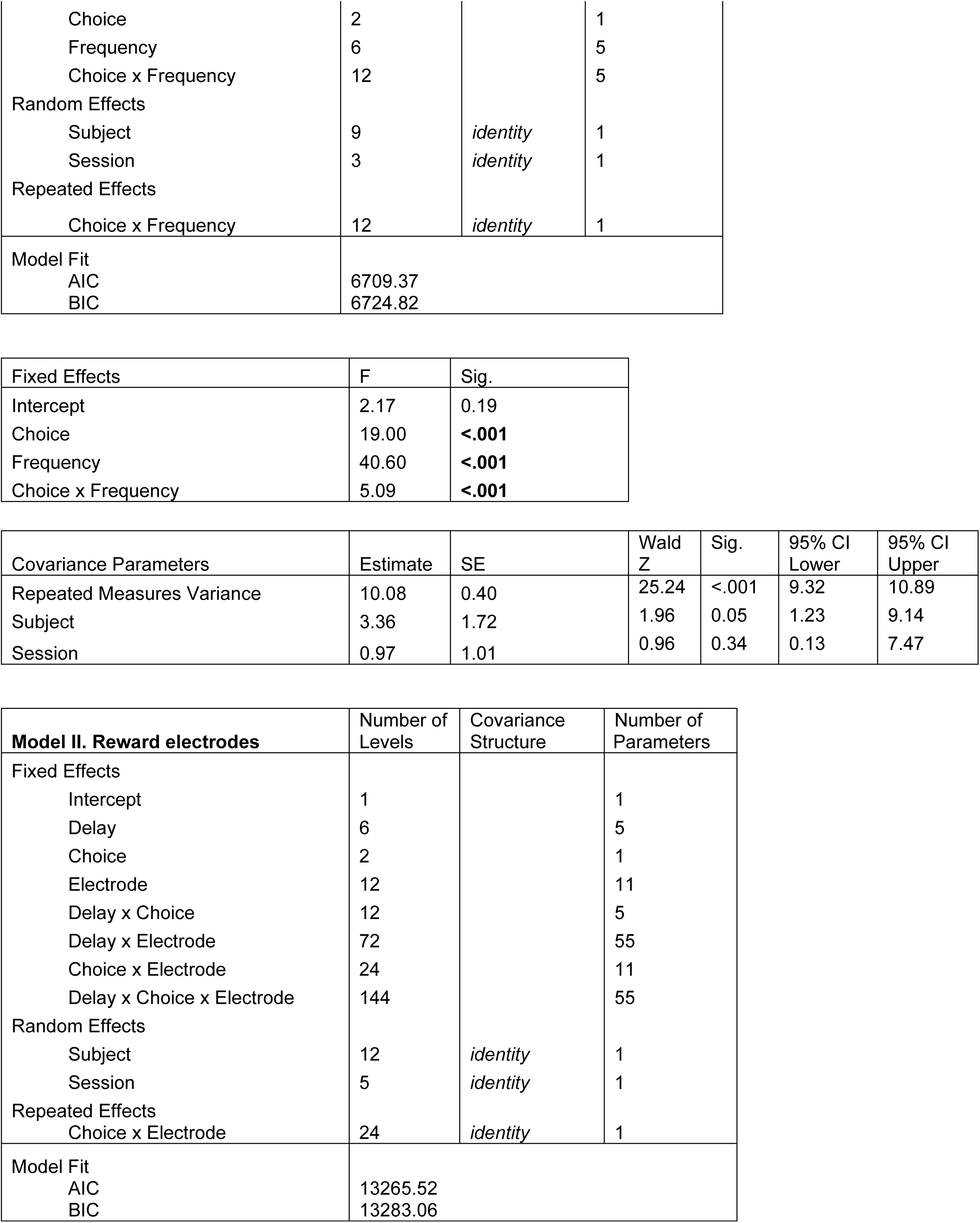
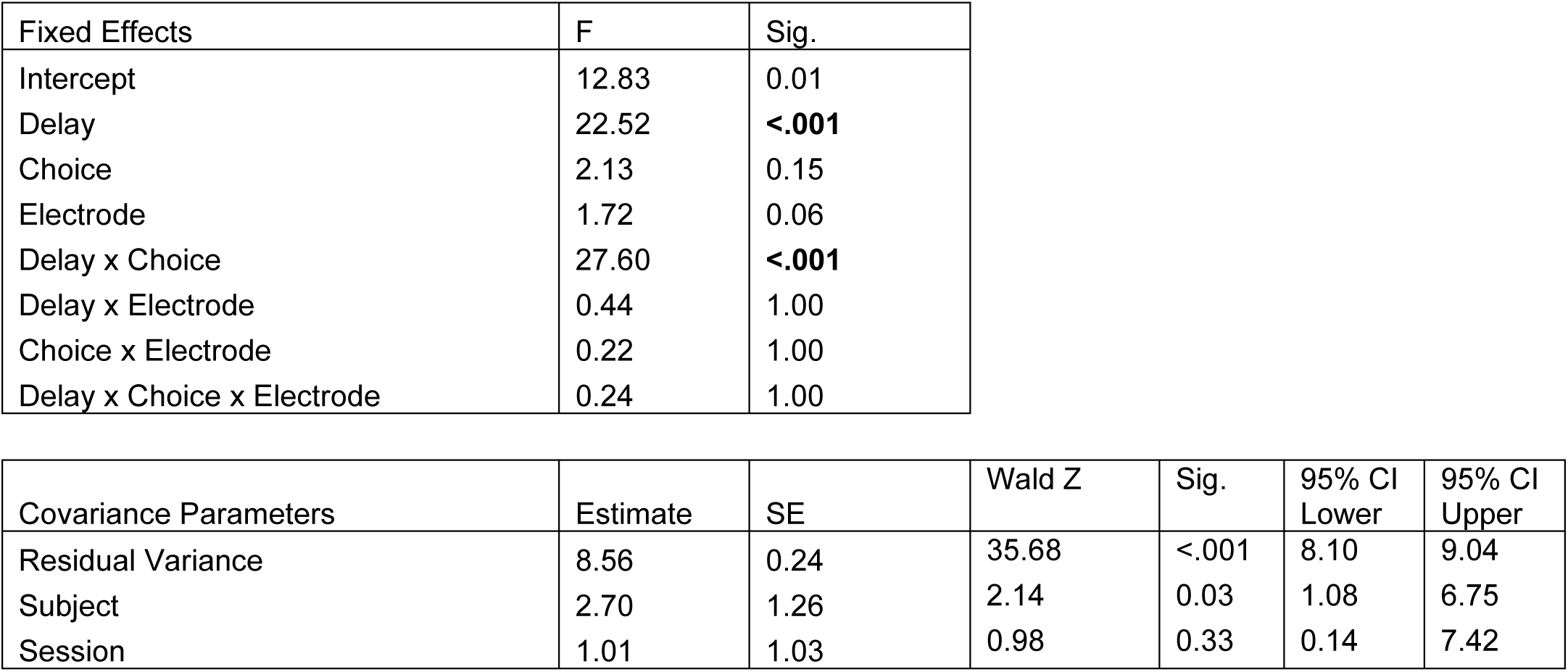
Linear mixed model design, fixed effects, and covariance parameter to explore power differences during reward outcome on the temporal discounting task.

| Model Dimensions |  |  |  |
| --- | --- | --- | --- |
| Model I. LOFC electrode | Number of Levels | Covariance Structure | Number of Parameters |
| Fixed Effects |  |  |  |
| Intercept | 1 |  | 1 |
| Choice | 2 |  | 1 |
| Frequency | 6 |  | 5 |
| Choice x Frequency | 12 |  | 5 |
| Random Effects |  |  |  |
| Subject | 9 | *identity* | 1 |
| Session | 3 | *identity* | 1 |
| Repeated Effects |  |  |  |
| Choice x Frequency | 12 | *identity* | 1 |
| Model Fit |  |  |  |
| AIC | 6709.37 |  |  |
| BIC | 6724.82 |  |  |

| Fixed Effects | F | Sig. |
| --- | --- | --- |
| Intercept | 2.17 | 0.19 |
| Choice | 19.00 | **<.001** |
| Frequency | 40.60 | **<.001** |
| Choice x Frequency | 5.09 | **<.001** |

| Covariance Parameters | Estimate | SE | Wald Z | Sig. | 95% CI Lower | 95% CI Upper |
| --- | --- | --- | --- | --- | --- | --- |
| Repeated Measures Variance | 10.08 | 0.40 | 25.24 | <.001 | 9.32 | 10.89 |
| Subject | 3.36 | 1.72 | 1.96 | 0.05 | 1.23 | 9.14 |
| Session | 0.97 | 1.01 | 0.96 | 0.34 | 0.13 | 7.47 |

| **Model II. Reward electrodes** | Number of Levels | Covariance Structure | Number of Parameters |
| --- | --- | --- | --- |
| Fixed Effects |  |  |  |
| Intercept | 1 |  | 1 |
| Delay | 6 |  | 5 |
| Choice | 2 |  | 1 |
| Electrode | 12 |  | 11 |
| Delay x Choice | 12 |  | 5 |
| Delay x Electrode | 72 |  | 55 |
| Choice x Electrode | 24 |  | 11 |
| Delay x Choice x Electrode | 144 |  | 55 |
| Random Effects |  |  |  |
| Subject | 12 | *identity* | 1 |
| Session | 5 | *identity* | 1 |
| Repeated Effects |  |  |  |
| Choice x Electrode | 24 | *identity* | 1 |
| Model Fit |  |  |  |
| AIC | 13265.52 |  |  |
| BIC | 13283.06 |  |  |

| Fixed Effects | F | Sig. |
| --- | --- | --- |
| Intercept | 12.83 | 0.01 |
| Delay | 22.52 | **<.001** |
| Choice | 2.13 | 0.15 |
| Electrode | 1.72 | 0.06 |
| Delay x Choice | 27.60 | **<.001** |
| Delay x Electrode | 0.44 | 1.00 |
| Choice x Electrode | 0.22 | 1.00 |
| Delay x Choice x Electrode | 0.24 | 1.00 |

| Covariance Parameters | Estimate | SE | Wald Z | Sig. | 95% CI Lower | 95% CI Upper |
| --- | --- | --- | --- | --- | --- | --- |
| Residual Variance | 8.56 | 0.24 | 35.68 | <.001 | 8.10 | 9.04 |
| Subject | 2.70 | 1.26 | 2.14 | 0.03 | 1.08 | 6.75 |
| Session | 1.01 | 1.03 | 0.98 | 0.33 | 0.14 | 7.42 |

When delays for each choice were equal (500ms), the linear mixed model revealed a main effect of choice (high vs. low reward) (F_(1,1274.92)_=19.00, *p <0.001*), a main effect of frequency (F _(5,1273.94)_ = 40.60, *p<0.001*) and an interaction between choice and frequency (F_(5,1273.94)_ =5.09; *p<.001*). Post-hoc tests (Bonferroni corrected) show the significant interaction was driven by a difference in power between high reward choice (EMM= 3.86, SE=0.89, CI= 1.83, 5.90) and low reward choice (EMM= 1.72, SE=0.89, CI =−0.87, 3.22) at beta and high-gamma (high reward choice; (EMM = 2.74, SE=0.89, CI= 0.70, 4.77) low reward choice (EMM= 2.16, SE=0.89, CI=0.11, 4.20) frequencies (**Fig. 3C**). Sessions contributed to only 6.7% of the total variance, but subjects contributed to 23% of the variance-a significant effect (Waldz 1.96, *p<.05*). Subjects did show individual differences in beta power values on high and low reward choices at matching delays (500ms) (**Supp Fig. 2**).

The next question we asked is whether reward-locked beta power was sensitive to the temporal delays of reward. If beta power reflected reward value, we hypothesized that the difference between high and low-rewards should be modulated with increasing delays, reflecting the discounted value of the delayed high-value choice. We used a second linear mixed model design to statistically measure whether there was an effect of delay length (0.5,1,2,5,10,20s), choice (high vs. low reward) and electrode location (M2, A32D, A32V, vOFC, ALM, LFC, Ains, LOFC, VMS, NAcS, NAcC, and BLA) on beta power (dependent variable) during reward feedback. Subject and session were used as random effects to observe their contribution to the model. We found a main effect of high-reward delay (F_(5,2533.84)_= 22.52, *p <0.001*) and an interaction between delay and choice (F_(5,2550.09)_ =27.60; *p<.001*). Post-hoc (Bonferroni corrected) tests performed for the lOFC electrode at each delay condition showed that at low delays (0.5s, 1s) there was greater beta activity for high-value choices (0.5s delay, estimated marginal mean (EMM) difference [high-low] in power = 2.69, SE of difference= 0.03; 1s delay, EMM difference = 1.02, SE of difference= 0.02). At a moderate delay (2s) there was no difference in power ([high-low] EMM =−0.06, SE of difference= 0.06); and with longer delays (5s, 10s, 20s) there was greater beta power on low-value choices (5s delay EMM [high-low] =−0.60, SE of difference= 0.02; 10s delay EMM difference =−1.33, SE of difference= 0.00; 20s delay EMM difference = −1.26, SE of difference= 0.08) (**Fig. 3D**). Thus, reward-locked power at beta frequencies in lOFC significantly decreases as value of high reward is less at larger temporal delays. Across the 12 putative reward regions (M2, A32D, A32V, vOFC, ALM, LFC, Ains, lOFC, VMS, NAcS, NAcC, BLA) there was no significant difference in beta power between electrode locations (*p<0.06*) (**Fig. 3E**). Thus, temporal discounting of beta power seems to reflect a value signal that is dispersed broadly across areas of the cortico-striatal reward network. Each subject had a slightly different beta power discounting curve (**Supp Fig. 2**) shown at the lOFC electrode.

Finally, to understand whether beta-oscillatory activity within this reward network was related to behavioral choice (i.e., preference for selecting either the high or low-value choice), we performed a logistic regression analysis with mean beta frequency power on a particular session as the dependent variable and the overall likelihood of choosing the high-value choice in that session as the independent variable. A positive beta value indicated a significant relationship between relative difference in beta power and the percent of high-value choices. We ran this analysis for each of the 12 brain regions, followed by FDR correction, using power from the difference (high-low reward) between trials. All 12 brain areas showed significant (FDR-corrected) positive relationships with high-reward choice and the differential beta power from the high and low-value responses (**Fig. 3F**). This suggests that the relative difference in beta power between high and low reward reflects value-related value that is directly linked, on a session by session basis with the choice animals make.

### Beta Power Reflects Reward Certainty and Updates after Reversal

On a probabilistic reversal learning (PRL) task, subjects first learned that one response leads to a high-probability of reward (“target”) and an alternate response would lead to a low-probability of reward (“non-target”), then subjects flexibly updated this representation after contingencies were reversed. In our version of this task, each day one response port would be randomly assigned to start as the target NP (rewards delivered 80% of the time) while the other, non-target, port would deliver rewards 20% of the time and reverse when 8 out of the last 10 trials were target choices (regardless of reward outcome) (**Fig. 4A**). Thus, on any session, subjects needed to dynamically modulate their behavior to track reward contingencies. The following analyses are from 79 behavioral sessions, from 7 male rats. The minimum number of behavioral sessions/ rat was 7. 36 PRL sessions included LFP data (average= 5 LFP sessions/ rat) (**Table 1**). On the first session rats performed an average of 1.33 reversals (SEM=0.211) and each animal showed significant improvement in number of reversals across time (t_(6)_ = 4.39, *p=.007,* paired t-test) (**Fig. 4B**). On the last session rats performed an average of 10.7 reversals (SEM=2.03). Rats tended to make the same choice after receiving a reward (termed a “win-stay” response). On 64.2 +/− 10.9% of rewarded trials, rats returned to the same response port on the subsequent trial (**Fig. 4B**).

**Figure 4:**
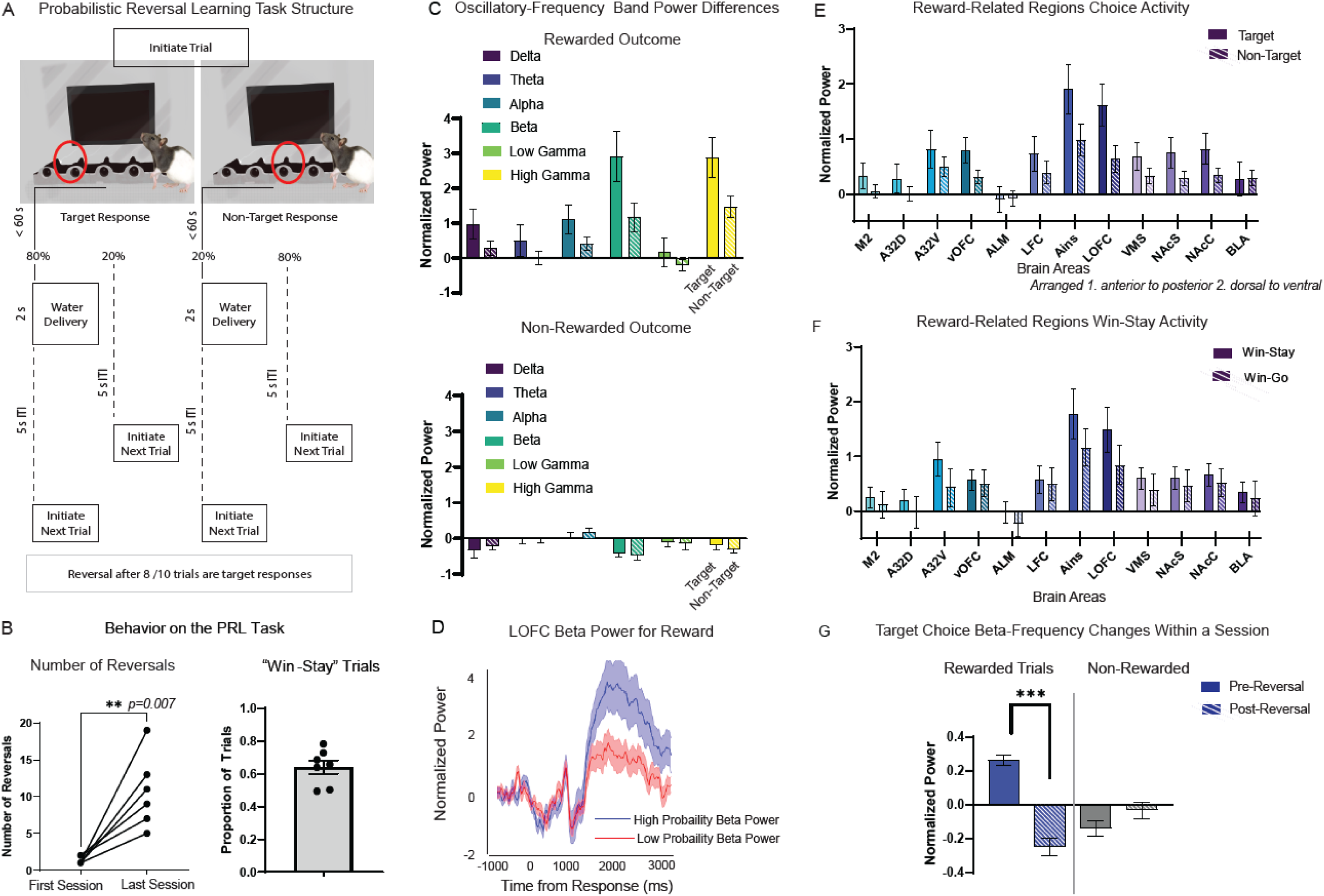
Likelihood of Reward Outcome Represented on the PRL Task. (A) Trial structure of the probabilistic reversal learning (PRL) task. Rats choose between two response ports: the target choice delivered reward 80% of the time and the non-target choice delivered reward 20% of the time. Reward contingencies reversed after 8 out of the last 10 trials were target choices (regardless of reward outcome). (B) Behavior on the PRL task measured by number of reversals per session and proportion of win-stay trials. The number of reversals is compared in the first and last session of individual rats to show improvement over time. The average and SEM proportion of win-stay trials/ session is plotted with black dots visualizing the average for each individual rat. (C) Average target (solid) and non-target (striped) choice power across delta (1-4 Hz), theta (4-8 Hz), alpha (8-12 Hz), beta (15-30 Hz), low gamma (50-70 Hz) and high gamma (70-150Hz) frequencies on the lOFC electrode during reward-feedback (500-2500ms) plotted separately for rewarded or non-rewarded outcomes. (D) The shaded error bar plot shows mean and SEM traces of lOFC normalized beta power for high-probability rewards (blue) and low-probability rewards (red). Traces are time-locked to response. (E) Beta power during reward-feedback (averaged activity from 5000ms-2500ms after response) across the 12 brain regions. The average and SEM are shown separately for target (solid) and non-target (striped) rewarded choices in each brain region. Brain regions are organized from 1. anterior to posterior and 2. dorsal to ventral. (F) Similarly, we also show the average and SEM beta power activity on win-stay (solid) compared to win-go (striped) trials in the 12 brain regions. (H) Beta power (from all 12 brain regions) is compared before and after a reversal on target choices. The average and SEM are shown separately for rewarded trials pre-reversal (light blue), rewarded trials post-reversal (dark blue), non-rewarded trials pre-reversal (light orange) and non-rewarded trials post-reversal (dark orange).

Based on data gathered from the temporal discounting task, we hypothesized that beta power may reflect subjective value, dynamically adjusting according to the value representation within a specific context. Thus, we expected to see greater beta power on rewarded target choices (high reward probability), compared to rewarded non-target choices (low reward probability). We predicted that beta-oscillations would dynamically track the choice leading to the higher-expected value and shift power after a reversal. To statistically model the effects of choice on beta power, we used a linear mixed model to compare power (dependent variable) on the lOFC electrode during the reward outcome period with frequencies (delta, theta, alpha, beta, low-gamma, high-gamma), choice (target vs. non-target) and outcome (reward or no reward) (**Table 5**).

**Table 5:**
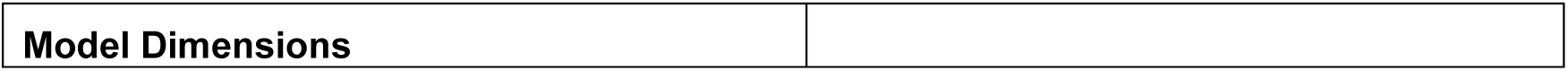
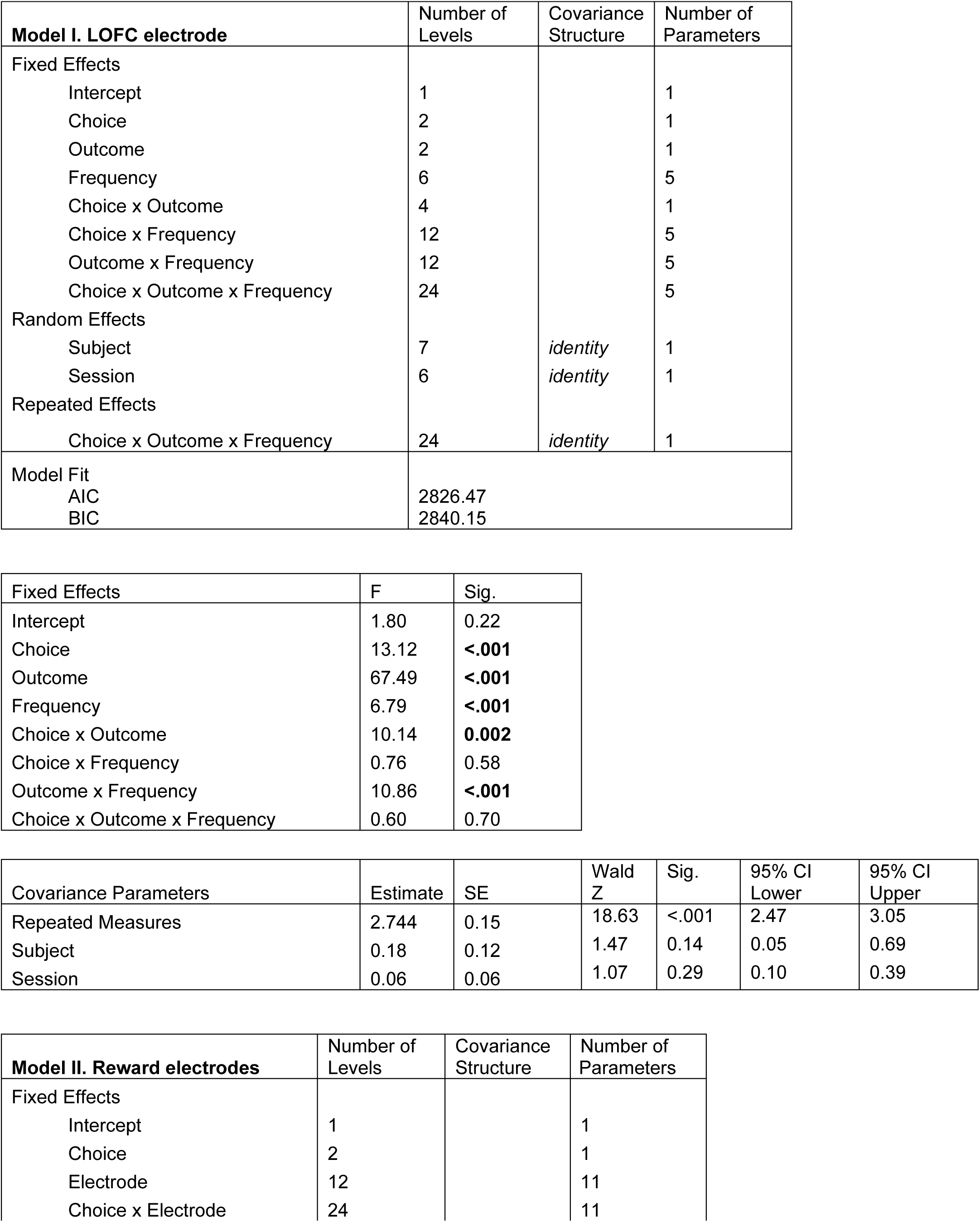
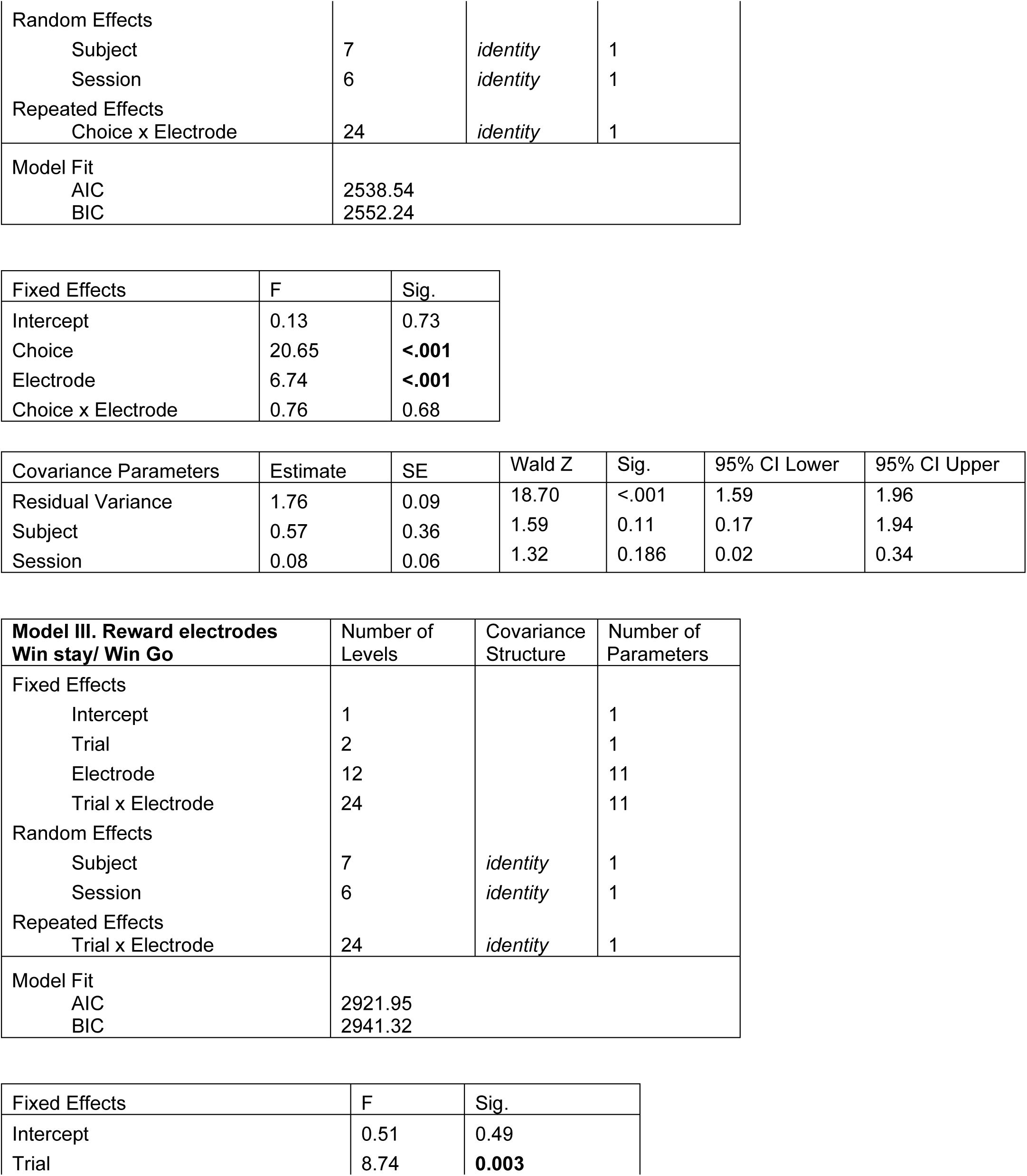
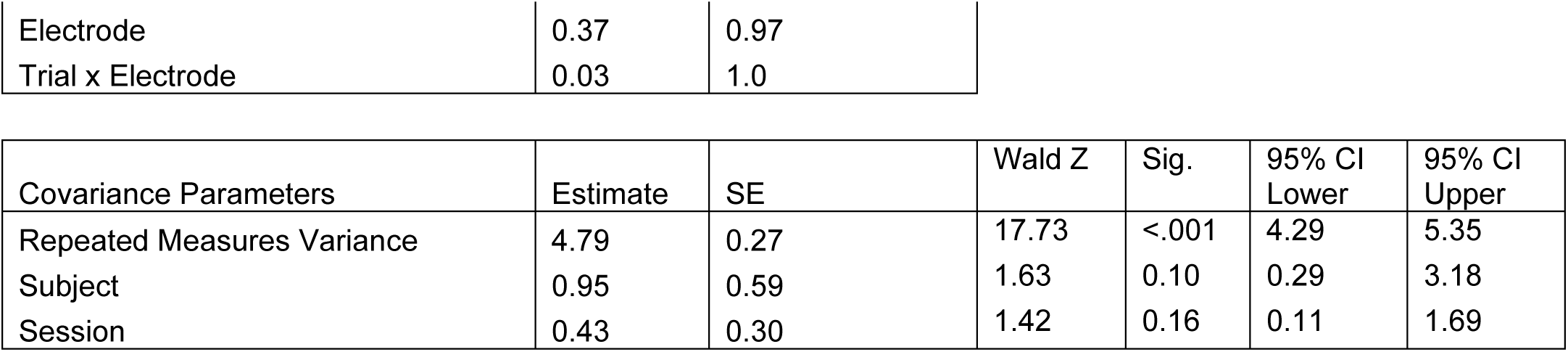
Linear mixed model design, fixed effects, and covariance parameter to explore power differences during reward outcome on the probabilistic reversal learning task.

|  |
| --- |
| **Model Dimensions** |

| **Model I. LOFC electrode** | Number of Levels | Covariance Structure | Number of Parameters |
| --- | --- | --- | --- |
| Fixed Effects |  |  |  |
| Intercept | 1 |  | 1 |
| Choice | 2 |  | 1 |
| Outcome | 2 |  | 1 |
| Frequency | 6 |  | 5 |
| Choice x Outcome | 4 |  | 1 |
| Choice x Frequency | 12 |  | 5 |
| Outcome x Frequency | 12 |  | 5 |
| Choice x Outcome x Frequency | 24 |  | 5 |
| Random Effects |  |  |  |
| Subject | 7 | *identity* | 1 |
| Session | 6 | *identity* | 1 |
| Repeated Effects |  |  |  |
| Choice x Outcome x Frequency | 24 | *identity* | 1 |
| Model Fit |  |  |  |
| AIC | 2826.47 |  |  |
| BIC | 2840.15 |  |  |

| Fixed Effects | F | Sig. |
| --- | --- | --- |
| Intercept | 1.80 | 0.22 |
| Choice | 13.12 | **<.001** |
| Outcome | 67.49 | **<.001** |
| Frequency | 6.79 | **<.001** |
| Choice x Outcome | 10.14 | **0.002** |
| Choice x Frequency | 0.76 | 0.58 |
| Outcome x Frequency | 10.86 | **<.001** |
| Choice x Outcome x Frequency | 0.60 | 0.70 |

| Covariance Parameters | Estimate | SE | Wald Z | Sig. | 95% CI Lower | 95% CI Upper |
| --- | --- | --- | --- | --- | --- | --- |
| Repeated Measures | 2.744 | 0.15 | 18.63 | <.001 | 2.47 | 3.05 |
| Subject | 0.18 | 0.12 | 1.47 | 0.14 | 0.05 | 0.69 |
| Session | 0.06 | 0.06 | 1.07 | 0.29 | 0.10 | 0.39 |

| **Model II. Reward electrodes** | Number of Levels | Covariance Structure | Number of Parameters |
| --- | --- | --- | --- |
| Fixed Effects |  |  |  |
| Intercept | 1 |  | 1 |
| Choice | 2 |  | 1 |
| Electrode | 12 |  | 11 |
| Choice x Electrode | 24 |  | 11 |
| Random Effects |  |  |  |
| Subject | 7 | *identity* | 1 |
| Session | 6 | *identity* | 1 |
| Repeated Effects |  |  |  |
| Choice x Electrode | 24 | *identity* | 1 |
| Model Fit |  |  |  |
| AIC | 2538.54 |  |  |
| BIC | 2552.24 |  |  |

| Fixed Effects | F | Sig. |
| --- | --- | --- |
| Intercept | 0.13 | 0.73 |
| Choice | 20.65 | **<.001** |
| Electrode | 6.74 | **<.001** |
| Choice x Electrode | 0.76 | 0.68 |

| Covariance Parameters | Estimate | SE | Wald Z | Sig. | 95% CI Lower | 95% CI Upper |
| --- | --- | --- | --- | --- | --- | --- |
| Residual Variance | 1.76 | 0.09 | 18.70 | <.001 | 1.59 | 1.96 |
| Subject | 0.57 | 0.36 | 1.59 | 0.11 | 0.17 | 1.94 |
| Session | 0.08 | 0.06 | 1.32 | 0.186 | 0.02 | 0.34 |

| **Model III. Reward electrodes**<br>**Win stay/ Win Go** | Number of<br>Levels | Covariance<br>Structure | Number of<br>Parameters |
| --- | --- | --- | --- |
| Fixed Effects |  |  |  |
| Intercept | 1 |  | 1 |
| Trial | 2 |  | 1 |
| Electrode | 12 |  | 11 |
| Trial x Electrode | 24 |  | 11 |
| Random Effects |  |  |  |
| Subject | 7 | *identity* | 1 |
| Session | 6 | *identity* | 1 |
| Repeated Effects |  |  |  |
| Trial x Electrode | 24 | *identity* | 1 |
| Model Fit |  |  |  |
| AIC | 2921.95 |  |  |
| BIC | 2941.32 |  |  |

| Fixed Effects | F | Sig. |
| --- | --- | --- |
| Intercept | 0.51 | 0.49 |
| Trial | 8.74 | **0.003** |

|  |  |  |
| --- | --- | --- |
| Electrode | 0.37 | 0.97 |
| Trial x Electrode | 0.03 | 1.0 |

| Covariance Parameters | Estimate | SE | Wald Z | Sig. | 95% CI Lower | 95% CI Upper |
| --- | --- | --- | --- | --- | --- | --- |
| Repeated Measures Variance | 4.79 | 0.27 | 17.73 | <.001 | 4.29 | 5.35 |
| Subject | 0.95 | 0.59 | 1.63 | 0.10 | 0.29 | 3.18 |
| Session | 0.43 | 0.30 | 1.42 | 0.16 | 0.11 | 1.69 |

Subject and session were investigated as random effects. On the lOFC electrode, there was a main effect of choice (F_(1,698.74)_= 13.12, *p<.001*), outcome (F_(1,697.98)_= 67.49, p<.001), and frequency (F_(5,694.38)_ =6.79, *p<.001*) and significant interactions between choice and outcome (F_(1,696.26)_ = 10.14, *p=0.002*) and outcome and frequency (F_(5,694.33)_ =10.86, *p<.001*) (**Fig. 4C**). Power was greater for rewarded (EMM=0.78, SEM= 0.21, CI= 0.30, 1.26) compared to non-rewarded outcomes (EMM= −0.24, SEM=0.21, CI= −0.72, 0.24) across all frequencies (main effect of outcome). There was also greater power for target choice (EMM=0.50, SEM=0.21, CI=0.02, 0.98) compared to non-target choice (EMM= 0.05, SEM= 0.21, CI= −0.43, 0.53) across all frequencies (main effect of choice). The interaction between outcome and choice showed beta and high gamma frequencies had the greatest power for target choice rewarded outcomes. Beta activity on the lOFC electrode during reward outcome was greater for target choice rewards (EMM= 2.39, SEM= 0.35, CI= 1.69, 3.09) compared to non-target choice rewards (EMM=0.90, SEM=.36, CI=0.19, 1.60). High-gamma power was also greater for target choice rewards (EMM= 2.69, SEM=0.35, CI= 1.99, 3.38) compared to non-target choice rewards (EMM=1.35, SEM=0.36, CI= 0.64, 2.06). Neither frequency showed significant differences in beta activity for non-rewarded target vs. non-target choices. (**Fig. 4C**). The shaded error plot illustrates the increased beta power during rewarded trials that is greater for high-probability (target) compared to low-probability (non-target) rewards at the lOFC electrode (**Fig. 4D**). Subject accounted for 6.0% of variance in our model and session accounted for 2.0%, neither which were not significant contributors to overall variance.

We followed this up with a linear mixed model to measure beta power (dependent variable) during rewarded outcomes across the other 12 reward-related electrodes for target and non-target choice (**Table 5**). There was a significant main effect of choice (F_(1,701.04)_ =20.65, *p<.001*) and a main effect of electrode (F_(11, 699.25)_ =6.74, *p<.001*), but no significant interaction between choice and electrode (F_(11, 699.21)_ =0.76, *p=.68*) (**Fig. 4E**). Post-hoc (Bonferroni corrected) tests revealed the main effect of electrode was influenced by increased power within anterior insula (EMM= 1.25, SEM=0.83, CI= −0.81, 3.30) and lOFC (EMM= 0.92, SEM= 0.83, CI= −1.14, 2.97) brain regions showing overall greater reward-related activity compared to others. Subjects contributed to 23.7% of the variance and session to 3.3%. Neither were significant contributors based on the Waldz test.

We next performed an analysis to see if beta-power on a trial was linked with activity on the subsequent (next) trial. Using a linear mixed model, we compared beta power for “win-stay” trials (rewarded trials in which animals chose the same response on a subsequent trial) and “win-go” trials (rewarded trials in which animals chose the different response on the subsequent trial) in all 12 electrodes. We observed a main effect of trial type (F_(1, 630.66)_ = 8.74; *p=<0.001*), and no significant interaction between trial type and electrode (F_(11, 628.38)_ =0.03, *p=0.97)* (**Fig. 4F**). Across electrodes, beta power was greater on win-stay trials (EMM=0.59, SEM=0.47, CI=−0.46, 1.63) than win-go trials (EMM= 0.08, SEM=0.48, CI −0.97, 1.13). Subject accounted for 15.4% of variance and session for 7.0%. Neither were significant contributors according to a Waldz test.

Finally, pooling data across all 12 brain regions, we examined how beta power reflected a change in reward contingencies. We used a two-way ANOVA to compare beta power on rewarded and non-rewarded target choices before and after a reversal. Specifically, we analyzed the last four “target” and “non-target” rewarded trials pre-reversal and the first four “target” and “non-target” rewarded trials post-reversal from the “new” target; and the same for non-rewarded trials. Analyzing the data this way we found a main effect of reward outcome (F_(1,332)_=12.0, *p*<0.001) on beta power in reward regions, no main effect of reversal (pre vs. post) (p=0.133), but a significant interaction between reward outcome and reversal (F_(1,332)_=13.4, *p*<0.001) (**Fig. 4G**). Post-hoc (Bonferroni corrected) comparisons show a selective decrease in beta power after a reversal that only occurs on rewarded target trials. The mean difference of beta power pre vs. post reversal was 0.521, SE of difference= 0.062, *corrected p <0.001* on rewarded trials. The mean difference of beta power pre-post reversal was −0.108, SE of difference= 0.062, *corrected p=0.166*, on non-rewarded trials (**Fig. 4G**). This is largely consistent with the idea that beta power reflects the expected outcome value which, immediately after a reversal, is still low for the “new” target. Beta-oscillations thus seem to reflect accumulating evidence about rewarded outcomes and modulating expectancy by tracking repeated positive outcomes and does not meaningfully reflect a signal related to the lack of reward (expected or unexpected).

### Verification of LFP Probe Locations at Target Brain Areas

Coronal sections stained with thionine to capture cell bodies were used to verify the electrode placement in target brain regions. For each cannula (1–8), a graphical representation of a rat brain atlas ((52) shows the identified center of recording sites at each DV location (four per cannula) (**Supp Fig. 3**). Colored dots represent the task the animals belong to (green: go/wait N= 6/11; pink: temporal discounting N= 9/10; blue: PRL N=7/7). An example coronal slice at the corresponding AP location is also shown for each cannula placement with magnification of each track in the brain. The table includes the AP, ML, and DV coordinates for all 32 electrodes and their corresponding nomenclature. The location of single-unit OFC recording electrodes is also shown (B) from a range of +4.2 AP through +3.25 AP relative to bregma. The LO/VO subdivisions are outlined on the example coronal sections taken from the rat brain atlas. Electrodes span both divisions. The graphical representation includes all electrode tracks (N=5/8 rats) (**Supp Fig. 3**).

## Discussion

Our results show changes in beta (15-30 Hz) and high-gamma (>70 Hz) frequencies that, across multiple distinct tasks scaled dynamically according to markers of learned and expected reward value. Each task contributed something unique to our findings and using different cohorts of animals offered replication for greater certainty of our findings. For example, measuring LFP on a behavioral inhibition task (go/wait), we identified beta power-related changes that signaled positive valence (rewarded) trials during reward-feedback. Firing rates of single-units in OFC were also modulated at beta frequencies during positive reward outcome; and the magnitude of the beta power was correlated with overall performance on that session, suggesting a relationship between beta power and reward expectation. On a temporal discounting task, beta power corresponded to subjective reward value and was significantly linked with choice for the immediate vs. the delayed condition. On a PRL task, beta was elevated for high-probability target choices; and higher beta power was associated with selecting the same trial following a rewarded outcome. We generally see evidence of a beta reward processing signal broadly throughout the cortico-striatal network, but there are instances where distinct brain regions are more/less engaged based on task-dependent features. A32D, lOFC, and anterior insula electrodes showed the most consistently elevated beta power across tasks during positive reward feedback, suggesting this effect was strongest in those cortical regions. Subtle variations in activity between tasks may represent examples of functional segregation between cortical subdivisions seen previously on other reward-guided tasks (9,29,41,53–55). For instance, ventral regions of striatum and orbitofrontal cortex show large increases in beta frequency power during rewarded outcomes only in the go/wait task where reward valance certain and less subjective. Moreover, elevated beta power may not always promote optimal behavior based on brain region and task-specific parameters. For instance, researchers using 20Hz (beta frequency) optogenetic stimulation of glutamatergic ventral medial OFC neurons found that activation impaired PRL performance whereas inhibition increased the number of reversals (56). We find a positive relationship between lOFC reward-locked beta power and behavior, but that relationship is likely different amongst cortical subregions. Thus, in the cortico-striatal network we find reward-locked oscillations at beta frequencies, in both single units and local field potentials, that mark positive reward valence and scale with reward expectation. Our findings are consistent across three different reward processing tasks suggesting that beta-oscillations may serve as a stable and robust bio-marker for future studies.

Data from each of these tasks, when considered alone, could have multiple explanations and confounds – however, the similar relationships between beta power and expected value observed across animals/tasks help define the role of beta-oscillations in reward processing. The most trivial explanation of our findings is that beta activity reflects a non-neural artifact time-locked with reward delivery such as movement (i.e. muscle/EMG-related contamination during reward consumption) or electrical noise associated with reward-delivery. However, it is not obvious how this explanation would show why on matched delays (500ms) on the temporal discounting task there was greater beta power for high-value compared to low-value trials during the first second following reward delivery when movements and electrical noise would at least in theory be matched. Similarly, data from the PRL task indicates beta power was greater for high-probability responses which also has identical reward delivery to low-probability responses.

A different possibility is that the beta activity is neuro-physiological in nature but reflects a motor, opposed to reward, process. Beta-oscillations have been well-characterized within motor cortex (57–61) and dorsolateral striatum (58,60,62,63) and tend to be largest pre/post-movement, but are classically reduced during movement (59,61,64,65). This functional description fits with observations of beta activity in Parkinson’s disease patients who have trouble initiating movements and show increased beta-oscillations related to symptom severity (59,62–64,66). Thus, one explanation is that increased beta power reflects motor inhibition that might occur and be linked with reward consumption. However, we believe our data is not compatible with this hypothesis in a few ways. First, sensorimotor beta-oscillations, as previously described, are more localized within motor and dorsal striatum, whereas we observe oscillations (and single-units related to beta-oscillations) more strongly within ventral brain regions (orbitofrontal cortex and insula, for example). Second, as before, we believe our data comparing high vs. low value reward (temporal discounting) and high-probability vs. low-probability reward (PRL) argues against this interpretation, as it is unclear why animals would be more stationary when consuming rewards on these trials where motor requirements should in theory be matched. It may be possible that animals more vigorously consume reward when there is a greater expectation of reward – in other words, that the neurophysiological processes we observe are, indeed, matched by a physical aspect. If this is the case, our results would still be valid, though the interpretation would be different. We do not currently have the data we need (high-frequency video of the licks) to distinguish this, and this will need to be clarified with further research.

If beta-oscillations reflect reward processing, then what specific aspect of reward might they represent? We hypothesize that beta-oscillations reflect activity within a cortico-striatal network that drives optimal decision-making based on expected reward-value. We provide evidence that beta-oscillations during reward feedback modulate activity based on task variables such as reward magnitude, temporal delay, and probability of reward. Growing research has identified beta-oscillations outside of sensorimotor networks related to attention (67, 68), top-down processing ((65, 69), working memory (67,70–72)and outcome evaluation (45, 73). Beta frequency impairments have been observed in cases of depression, bipolar disorder, schizophrenia, attention disorders, and addiction (opioid and alcohol) (74). Consistent with our findings, beta-oscillations during reward-feedback have previously been observed in humans and animals. EEG and MEG measures in humans find beta oscillations during positive-valence reward within frontal-striatal circuits that is sensitive to reward valence, magnitude, and predicts subsequent choice ((71,75–79). Similarly, increased beta power in cortico-striatal regions has been observed in rodents approaching reward locations (47, 80)that was modulated by reward magnitude ((80)and stabilized with task experience (28, 47). Recently, it was observed that during a reward discrimination task, increased beta power 100-200ms after reward feedback in the anterior cingulate cortex and nucleus accumbens of rodents that was correlated with response bias (81)Our work extends this prior data by conclusively demonstrating a relationship between beta power and reward expectation across multiple task contexts. It further suggests that beta-oscillations can be utilized as a cross-species translational marker of value estimation that is linked to reward-guided behavior and could be used to predict reward sensitivity, risk-taking behavior, and impulsivity.

The feedback-related negativity ERP signal classically observed in humans is thought to reflect dopamine transmission (81–83), but the signal gets more negative following positive reward valence (81); the opposite of our beta oscillatory signal. Dopamine activity is linked with both reward-prediction and reward-prediction errors (RPE) (10,16,78,84). Previous research in humans explored the possibility of frontal beta-oscillations as an RPE signal but found that stimuli signaling expected rewards elicited more beta power than unexpected rewards; the inverse of an RPE (78). Our results are consistent with an inverse correlation between beta activity and the dopaminergic RPE signal. First, on the go/wait task we find beta power signals positive reward outcomes and correlates with more accurate task performance, whereas dopamine transmission would be higher when rats are performing poorly (more unpredictable). In the temporal discounting paradigm dopaminergic activity is greater for longer delay periods (19) corresponding with the reduction in beta power we observed during longer delays. Finally, more dopamine is observed for unexpected rewards, but on the PRL task we see higher beta power on expected reward outcomes that rapidly decays after a reversal when expectancy signals are not defined. Moreover, we find single-units in OFC that are correlated with beta oscillations during reward delivery that may be consistent with reward prediction rather than an error in prediction (43). Increased beta oscillations in frontal cortex could therefore be a marker of a suppression in dopamine release. Strikingly, an inverse relationship between dopamine and beta-oscillations has been observed in motor cortex and dorsolateral striatum as well (10, 63). Therefore, a similar relationship between beta-oscillations and dopamine may exist within ventral striatum and prefrontal cortex. In this way, beta signals may represent a common modality of communication across distributed cortico-striatal networks: cortico-thalamic-basal ganglia pathways for motor controls (60, 66)and cortico-striatal-limbic pathways for reward processing (2,6,8,11,12,18,85) (Schultz et al., 2000; Dalley et al., 2004; Abler et al., 2006; Berridge and Kringelbach, 2008; Haber and Knutson, 2010; Chau et al., 2015). A common striatal-beta generating mechanism could explain how increases in attention, motor inhibition, and reward processing information are linked to beta-oscillations in distributed brain networks (59, 67), and suggests that, perhaps, dopamine influences this transmission similarly across these cortico-striatal networks.

We acknowledge there are many subsequent analyses to be completed for each task. Here, we present a comprehensive overview of reward processing activity in all tasks opposed to the fine intricacies of each which may be best explored by fitting reinforcement models to examine trial x trial decision making behavior and oscillatory activity Future analyses will investigate network-level connectivity to determine whether beta-oscillations originate in brain areas, like the striatum, or if they are an emergent property of cortico-striatal networks. Moreover, investigations will need to extend beyond power measures to include phase dynamics which can determine temporal relationships between brain areas. Across our tasks, we also see evidence of increased high-gamma power during reward-feedback. Much like theta-gamma coupling is linked to learning and memory in rodents (87–89), beta-high-gamma coupling may be linked to reward processing or reflect spike coherence. Researchers have described beta/gamma event-related synchronization that occurs after reward feedback in lateral prefrontal cortex of humans (71, 78). Additionally, our results are limited to only male rats. We are now repeating this set of studies in a balanced cohort of male/females to understand whether these findings generalize across sexes. Finally, further analysis of movements/video-tracking would lend greater certainty to our findings and rule out movement-related artifacts. Based on the preponderance of evidence across animals and tasks, we propose that beta-oscillations during reward-feedback may present a phenotype that can be used to identify disturbed reward-related processing deficits in psychiatric disorders or brain injury.

## Material and Methods

### Ethics Statement

This research was conducted in strict accordance with the Guide for the Care and Use of Laboratory Animals of the National Institutes of Health. The protocol was approved by the San Diego VA Medical Center Institutional Animal Care and Use Committee (IACUC, Protocol Number A17-014).

### Experimental Design

#### Subjects

37 male Long-Evans rats obtained from Charles River Laboratories were used for these experiments. When received, rats were ~ one month old weighing 150g. Habituation and pre-training was initiated two weeks after arrival. Depending on the task, rats trained for 5-14 weeks before receiving surgery. Rats were housed in pairs during prior to electrode implantation, and individually housed thereafter, in a standard rat cage (10 x 10.75 x 19.5 in, Allentown, NJ, USA) with free access to food and on a standard light cycle (lights on at 6 am / off at 6 pm). During behavioral training, animals underwent water scheduling (free access to water for two hours/day) to maintain motivation for water reward in the tasks. Water was unrestricted on non-training days and rats were weighed weekly to ensure that water scheduling did not lead to reduced food intake. Different cohorts of rats were trained to perform one of three tasks designed to measure distinct aspects of reward processing12 rats with multi-site LFP probes were trained on a go/wait response inhibition task, 10 on a temporal discounting task, and 7 on a probabilistic reversal learning (PRL) task (**Table 1**). Additionally, 8 rats trained on the go/wait task were used for single-unit recordings to supplement LFP findings (**Table 1**). Subjects with chronic implants were monitored daily for signs of infections, injuries, and bleeding.

#### Operant Chamber and Training

The same custom-designed operant chamber was used for all three tasks. The chamber had five nose-ports (NP), each with an LED, IR sensor and metal cannula for water delivery. The chamber also contained two auditory tone generators, a house-light, a screen to display visual stimuli, and five peristaltic stepper motors/water pumps that delivered the water rewards into NPs. The chamber was 6.2 x4.7x 6.23 inches with a ceiling opening that allowed electrophysiology tethers to move freely. Simulink (Mathworks) installed directly onto a Raspberry Pi system controlled the behavioral programs. Behavioral outputs from the operant chambers were synchronized with electrophysiological signals using lab-streaming-layer, a protocol designed to integrate multiple behavioral and physiological streams into a common timing stream (90, 91). The design, operation and software control of this chamber has been described previously (90). Animals first went through a pre-training period (5-10 sessions), to learn that a NP with an LED “on” signaled an available response port; that responding in an available NP would trigger a water reward; and finally that there was a sequential nature to the task (animals start a “trial” by first entering the middle NP (3), after which they could use either of the neighboring ports (2 or 4) to respond and collect an immediate reward). This standard pre-training paradigm was used for all three final behavioral paradigms. Animals advanced to the next stage of training when they consistently performed ≥100 trials in a 60 min session.

#### Behavioral Tasks

*I. Go/Wait Response Inhibition Task.* The visual-cue go/wait task was used to observe brain activity associated with positive valence on successful go-cue trials (rewarded) compared to unsuccessful wait-cue trials (not rewarded). Animals began a trial by entering the middle NP (3), ensuring animals were in an identical position on every trial when the visual stimulus appeared. After a fixed delay of 30ms, a visual stimulus appeared on the screen denoting the trial as either a “go-cue” trial (animal required to respond within 2s to attain a reward) or a “wait-cue” trial (animal required to withhold response for 2s to attain a reward). The stimulus remained on the screen until the animal responded. If animals responded correctly, a water reward was delivered into the middle NP (3) after a delay of 400ms. If animals responded incorrectly, the house light flashed for a 5 second “time-out” period and no reward was given. Rewards consisted of 20 µL of water delivered over a two second period using a stepper-motor (the motor sound provides an instantaneous cue regarding reward delivery). After water delivery, there was a 5s inter-trial-interval before the next trial began. The trials were distributed randomly as 25% “go-cue” and 75% “wait-cue” trials. Animals were trained for ~14 weeks until behavior typically stabilized (>80% accuracy on go-cue trials), after which they were implanted with electrodes (described below). We waited two weeks for animals to recover from surgery prior to resuming water-scheduling. LFP analyses are based on data from 67 recording sessions from 12 rats (**Table 1**). Single-unit analyses are based on data from 62 recording sessions from 8 rats (**Table 1**).

*II. Temporal Discounting Task.* A different cohort of animals were trained on a temporal discounting task to contrast electrophysiology activity at different reward magnitude choices (high vs. low reward) delivered at increasing temporal delays (0.5 to 20s). Generally, temporal discounting tasks center around choosing between a low-value reward delivered immediately, or a high-value reward delivered after a delay. In our version of the task, subjects chose between a low-value (1x) reward delivered immediately (500ms after response) or a high-value (3x) reward delivered at a variable delay. In separate behavioral sessions the high-value delay ranged from 500ms to 20s after the response. Each session began with 6 forced-choice trials, orienting the rat to both the low-value (NP 2) and high-value (NP 4) options. The houselights were on, and LED lights signaled the available response port, alternating between low-value (NP 2) response and high-value (NP 4) response. Reward following either response was delivered immediately (500ms) after response during forced-choice trials. After the 6 trials were complete, the houselights dimmed and rats began the full, self-paced, trial sequence. Response port (2 and 4) LEDs were on, signaling the rat to choose. Selecting the low-value response port (2) turned on the houselights, the middle NP LED (3), and a tone (500ms duration) to indicate a choice was made. A small reward (10µL delivered over a 1s duration) was delivered immediately (500ms after response) from NP 3. Selecting the high-value response port turned on the houselights, the middle NP LED (3), and a tone (500ms duration) signaled the choice. Between sessions there was a variable delay (0.5s, 1s, 2s, 5s, 10s, 20s) until the high-value reward (30µL over a 3s duration) was delivered out of NP 3. The motor delivering water made an audible sound, to cue reinforcement delivery onset and amount of reward. The high-value delay alternated between behavioral sessions but remained the same throughout the entire (60 min) session. The houselights turned off when water was delivered out of NP3 and a 5s inter-trial interval began after water delivery. To learn to discriminate magnitude differences, rats were trained on the immediate delay condition (500ms) for both high-value and low-value choices. Once they showed a clear preference for the high-value choice (≥70% high-value responses/session) and consistently performed ≥100 trials, they were advanced to the other delay conditions. Training (including pre-training sessions) on average lasted 18 sessions across 5 weeks. After implantation we waited two weeks to allow animals to recover from surgery before electrophysiology recording began. Recording sessions lasted 60 minutes and occurred 3-4 days a week. LFP analyses are based on data from 124 recording session from 10 rats (**Table 1**). There was an average of 20 recording sessions per delay condition (two sessions at each delay condition per rat; N=19 sessions at 0.5s; N=23 sessions at 1s; N=15 sessions at 2s; N=28 sessions at 5s; N=18 sessions at 10s; N=21 sessions at 20s).

*III. Probabilistic Reversal Learning Task.* The probabilistic reversal learning task (PRL) was used to examine brain activity associated with learned reward likelihood (high vs. low-probability choice) and tests the subjects’ ability to update information after reward contingencies are reversed. In our version of the PRL task, rats must choose between two nose ports: the high-probability choice (“target“) delivers water 80% of the time, and the low-probability choice (“non-target”) only 20% of the time. The PRL task is self-paced. Each trial began with houselights off and the middle NP LED (3) on. Once a rat responded, LEDs in NP 2 and 4 turned on, indicating an available choice between the response ports. Each NP is randomly assigned as the target or non-target NP in each session. Selecting the target choice led to 2s (20µL) of water on 80% of trials and no water only 20% of the time. Selecting the non-target choice led to 2s (20µL) of water only 20% and no water 80% of the time. A response in NP 2 or 4 caused the other LED to turn off. On rewarded trials, the houselights remained off and water was delivered out of the selected NP (2 or 4) 500ms after the response. There was a 5s inter-trial interval that started with water delivery. On unrewarded trials, a tone (500ms in duration) signaled no water delivery, the houselights turned on, and a 5s inter-trial interval began. Throughout a session, the NP contingencies reversed based on the rats’ behavior. Reversals occurred when a rat made 80% target responses (rewarded or non-rewarded) over a 10-trial moving window (8 of the last 10 responses are “targets”). To perform PRL effectively, rats must respond appropriately to correct feedback (“target” rewards; “non-target” no reward) while also ignoring misleading feedback due to the probabilistic nature of our task (“target” no reward; “non-target” reward). Rats received at least two weeks of pre-training (described above) prior to surgical implantation of LFP probes but were naïve to the PRL task. Two weeks after surgery rats began training on the PRL task. Performance was measured by counting the number of reversals/ sessions, the target choice percentage, and win-stay behavior (propensity to choose the same NP after receiving a reward on the previous trial). On average, rats took 15 sessions to train on the PRL task (~3.5 weeks). Once rats were consistently performing at least one reversal and performing ≥100 trials we started to record LFP. Rats ran 60 min sessions 3-4 days a week. Behavioral data was collected from 7 rats across 79 PRL sessions, 36 of which included LFP recording (average of 5 sessions per rat) (**Table 1**).

### Surgery

Aseptic surgeries were performed under isoflurane anesthesia (SomnoSuite, Kent Scientific, CT, USA with all instruments autoclaved prior to start. Animals received a single dose of Atropine (0.05 mg/kg) to diminish respiratory secretions during surgery, a single dose of Dexamethasone (0.5 mg/kg) to decrease inflammation, and 1mL of 0.9% sodium chloride solution prior to surgery. The area of incision was cleaned with 70% ethanol and iodine solution. A local anesthetic, Lidocaine (max .2cc), was injected under the skin at the incision site while the animal was anesthetized but before surgery initiation. The fabrication and implantation procedures of our custom fixed field potential and single-unit probes are described in detail (37).

#### LFP Probe Implantation

Briefly, our LFP probe targets 32 different brain areas simultaneously. 50µm tungsten wire (California Fine Wire, CA, USA) used for our electrodes was housed in 30-gauge stainless steel metal cannula (Mcmaster-Carr, Elmhurst, IL, USA) cut 8-9mm long. Each cannula (N=8) contained four electrode wires cut to their unique D/V length. The average impedance of our blunt-cut tungsten microwires is 50 kOhms at 1 kHz. During surgery, 8 holes were drilled in the skull (one for each cannula) at predetermined stereotactic locations (see **Supp Fig. 3**). Additional holes were drilled for a ground wire and anchor screws (3–8). The ground wire was soldered to an anchor screw and inserted above cerebellum. Electrodes were slowly lowered to desired depth, pinned to the EIB board, and secured with superglue followed by Metabond (Parkell, NY, USA). The entire head stage apparatus was held to the skull and encased with dental cement (Stoelting, IL, USA).

#### Single-unit Probe Implantation

To record single-units we used a 32-channel stationary array with microwires arranged in a brush-like formation (see (37). Initial preparation of the animal and location of ground screws was identical to the LFP probe surgical procedures described above. A cranial window with diameter of 2mm was drilled with a 0.7mm micro drill (Stoelting, IL, USA) centered at the OFC target location. Three probes targeted ventral OFC (AP:+3.5mm, ML: +/−1.5mm, DV: 5.0mm), four targeted lateral OFC (AP: +3.5mm, ML:+/−2.5mm, DV: 5.0mm), and one implant had 16 electrodes in each region (**Supp Fig. 3**). The implant was lowered to desired depth slowly under stereotactic control. A thin layer of superglue was applied to the skull followed by a layer of Metabond (Parkell Inc., NY, USA) to seal the craniotomy. The implant was secured to anchor screws and attached to the dry skull with dental cement (Stoelting, IL, USA). The ground wire was pinned to the channel on the EIB board, and the remaining exposed wires covered in dental cement.

At the conclusion of surgery, the skin was sutured closed, and rats were given a single dose (1mg/kg) of buprenorphine SR for pain management. Rats recovered from surgery on a heating pad to control body temperature and received sulfamethoxazole and trimethoprim in their drinking water (60mg/kg per day for 8 days) to prevent infections.

### Electrophysiology

LFP data was recorded using a 32-channel RHD headstage (Intan Technologies, CA, USA; Part C3324) coupled to a RHD USB interface board (Intantech, Part C3100) and SPI interface cable. We used plug-in GUI (Open Ephys) software for acquisition. Data was recorded at 1Khz, with a band-pass filter set at 0.3 to 999 Hz during acquisition. Physiology data was integrated with behavioral data using a lab-streaming-layer (LSL) protocol (Ojeda et al., 2014), as described previously (90).

Single-unit data was recorded using a 32-channel RHD headstage with signal amplified using a PZ5 Neurodigitizer and RZ2 bioamp processor (TDT, FL, USA). Recorded signals were processed using Synapse software (TDT) at a sampling rate of 25KHz, high-pass filter of 300Hz and low-pass filter of 3000Hz. Behavioral markers were also integrated using LSL protocol.

### Statistical Analysis

#### LFP Time Frequency Analysis

We carried out standard pre-processing and time frequency (TF) analyses using custom MATLAB scripts and functions from EEGLAB (37–39). 1) Data epoching: We first extracted time-points for events of interest during each task. This paper focuses on neural activity time-locked to response/reward (opposed to trial onset) to examine neural activity during the reward-feedback period. Time-series data was extracted for each electrode (32), for each trial and organized into a 3D matrix (electrodes, times, trials). 2) Artifact removal: Activity was averaged across the time/electrodes to get a single value for each trial. Trials with activity greater than 4X standard deviation were treated as artifact and discarded. 3) Median reference: At each time-point, the “median” activity was calculated across all electrodes (32) and subtracted from each electrode as a reference. 4) Time-Frequency Decomposition: A trial by trial time-frequency decomposition (TF decomposition) was calculated using a complex wavelet function implemented within EEGLAB (newtimef function, using Morlet wavelets, with cycles parameter set to capture frequency windows of between 2 to 150 Hz (2 to 70 Hz in the go/wait task) and otherwise default settings used. We calculated the analytic amplitude of the signal (using the abs function in MATLAB). 5) Baseline normalization: To measure evoked activity (i.e. change from baseline) we subtracted, for each electrode at each frequency, the mean activity within a baseline window (1000-750ms prior to the start of the trial). 6) Trial averaging: We next calculated the average activity across trials for specific trial types at each time-point and frequency for each electrode, thus creating a 3D matrix (time, frequency, and electrode) for each behavioral session. Trials of interest were different for each task: In the go/wait task we analyzed go-cue rewarded trials, wait-cue rewarded trials and wait-cue unrewarded trials (go-cue unrewarded trials were too few to include); temporal discounting task we separated high and low-reward choice at each delay condition; and PRL task we separately analyzed high-probability (target) vs. low-probability (non-target) choices and their reward outcomes. 8) Comparison across animals: Prior to averaging across sessions/animals, we “z-scored” the data recorded from each behavioral session by subtracting the mean and dividing by the standard deviation of activity in each electrode (at each frequency) over time. Z-scoring was helpful for normalizing activity measured from different animals prior to statistical analysis. Because we had already performed a “baseline” subtraction (as described above), this analysis captured whether there was a significant increase or decrease in activity compared to baseline. FDR-correction was performed across all time-frequency-electrodes (FDR-corrected p-value threshold set to 0.05). These pre-processing steps resulted, for each session, in a 3D time-frequency-electrode matrix of dimensions 200×139×32 which was used for further statistical analyses as described below.

#### LFP Linear Mixed Models

We analyzed the time-frequency-electrode (TFE) data at the level of each session using linear mixed models in IBM SPSS Statistics v.28 (New York, United States) to account for subject and session variance. Across all three tasks we first used a LMM to compare normalized power (dependent variable) at each oscillatory frequency band in the LOFC electrode at trial types of interest. We used the following frequency bands: delta power (1-4 Hz), theta (4-8 Hz), alpha (8-12 Hz), beta (15-30 Hz), gamma (40-70Hz) and high gamma (70-150 Hz). Next, we used different LMMs to explore power (dependent variable) across 12 electrodes (M2, A32D, A32V, vOFC, ALM, LFC, Ains, lOFC, VMS, NAcS, NAcC, BLA) at time points of interest. Each model’s parameters including fixed, random, and repeated effects are specified for each analysis (**Table 3–5**). Data from the go/wait and PRL tasks was time-locked to response. We analyzed the full two second window of reward-feedback (500-2500ms after response). To account for the variable delay-to-reward conditions in the temporal discounting task, data was time-locked to reward delivery. We analyzed the first second of activity post-reward (0ms to 1000ms after reward onset) to control for the difference in water delivery between the high (3s) and low (1s) reward magnitudes.

We compared the Akaike information criteria (AIC) and Bayesian information criterion (BIC) of four commonly used covariance models (compound symmetry, scaled identity, AR(1), and unstructured) to determine the best fit (92, 93)The scaled identity model, assuming repeated measures may be independent but with equal variance (92, 93), provided the lowest AIC and BIC scores. We used a Restricted Maximum Likelihood (REML) model with the Satterthwaite approximation in SPSS. The fixed effects and estimates of each covariance parameter are reported for each test. Significant interactions with followed up with pairwise comparisons (Bonferroni corrected) in SPSS. Main effects of the Estimated Marginal Means of factors and their interactions were Bonferroni corrected. Linear Mixed Models account for missing data which was present in the subsequent analyses. For instance, the total number reported may be less than 12 brain areas x 128 sessions because in some sessions a particular electrode may not have provided usable data (noise/ broken channels, etc.).

### LFP Related to Behavioral Performance

In the go/wait task, the significant oscillatory frequencies (identified with the linear mixed model and post-hoc tests) were correlated with choice accuracy (in MATLAB). We correlated power on the lOFC electrode during wait-cue correct (rewarded) wait-cue incorrect (non-rewarded) and the difference in power between trial types with accuracy on wait-cue trials during the reward-outcome period (from 500 to 2500ms after response). We calculated correlations across all sessions/animals, followed by FDR correction.

In the temporal discounting task, we used regression models with the general linear model framework (in MATLAB) to compare the mean oscillatory frequency power (from 0 to 1000ms after reward) on a particular session was the dependent variable and the overall likelihood of choosing the high-reward choice on that session was the independent variable. We did not control for delay condition as we already determined in subsequent analyses that it was indeed a significant factor in modulating reward-related activity. We calculated correlations across all sessions/ animals for each of the 12 reward-related brain regions, followed by FDR correction.

In the PRL task, we used a two-way ANOVA to determine how significant oscillatory activity (defined in the linear mixed model) updated with reward contingency reversals. In a single session, we calculated the average power for the first four trials before a reversal and four trials following a reversal across all 12 electrodes during the reward outcome period (500-2500ms after response). Average activity before and after reversal was calculated separately at each electrode for each reversal pooled across animals/sessions.

#### Single-unit Analyses

Single-unit activity was recorded in 8 animals performing the go/wait task. We extracted behavioral markers of interest, LFP streams, and spiking data. Neural data was cleaned and referenced off-line using Wave_Clus v.2.5., a Matlab-based spike-sorting program (94). Signals were processed as two polytrode (16-electrode) groups with median referencing applied to each channel (see (37)). A threshold of spike detection was set at 5 times standard deviation of voltage potential for each channel. Broken channels, with large impedances beyond 10 MOhm were excluded from referencing/ clustering. Spikes in each polytrode group, as identified by Wave_Clus, were examined manually for characteristics of single-units (average spiking rate within the whole session was more than 0.5Hz, fewer than 1.5% inter-spike interval violations (<3ms), waveform resembling action potentials as opposed to sinusoidal noise artifacts, and clusters distinct from others in the principal component space) (95–97). Spikes meeting criteria were time-locked with behavioral events in MATLAB.

MATLAB functions were used to create peri-stimulus time histograms and raster plots at trial types of interest (go-cue rewarded / wait-cue unrewarded trials) to compare reward valence. Activity was time-locked to response (time point 0). PSTH (spikes/second) were made in 25ms bins from −2s to +2s after response and gaussian filtered for smoothing. Activity was baseline normalized by subtracting the average firing rate during the pre-response baseline (first 750ms from trial onset) on go-cue correct trials from firing rate in subsequent time bins. Units from recording days with at least 30 trials and a minimum baseline firing rate of 2 spikes/s were further analyzed for their task-related activity. A unit was considered “task-modulated” if the average firing rate was 2 standard deviations above or below the baseline activity for > 75 consecutive ms in either go-cued correct or wait-cued incorrect trials. Task-modulated units were categorized further based on the timing of their activation/ suppression relative to response. Anything with significant (+/− 2 standard deviations) activation or suppression from −500 to 0ms from response was considered pre-response activity (action). Anything with later activation or suppression (0 to 2000ms) was considered post-response (outcome) activity.

Finally, we calculated spike-field coherence (SFC) to relate spiking activity to field-potential oscillations. SFC ranges from 0 (spikes operate independently from LFP) to 1 (spikes are completely aligned to LFP) (44,46,48). To calculate SFC, we matched spike times to the continuous LFP signal from an electrode with minimal artifacts by multiplying the sampling rate of the LFP (1.0173). Next, we down sampled to normalize non-standard sampling frequencies and epoched the LFP data to align with task events (described above). SFC was analyzed separately at each oscillatory frequency (delta, theta, alpha, beta, gamma) for each trial type (go-cue correct and wait-cue incorrect), time-locked to response. Units were categorized as “high” or “low” SFC by using a median split based on beta frequency SFC values from 500-2500ms after response (reward feedback) on go-cued correct trials. Units one standard deviation above the median SFC value were categorized as “high” and units one standard deviation below were categorized as “low”. A two-way ANOVA was used to compare firing rate on go-cue correct vs. wait-cue incorrect trials x SFC value at beta frequencies (classified as “high” or “low”).

### Histology

Histological analyses were completed for 22/ 28 rats with LFP implants (go/wait task N=6/11; temporal discounting N=9/10; probabilistic reversal learning N=7/7) and for 5/8 rats with single-unit implants. At completion of recording sessions wire tips were marked by passing 12μA current for 10s through each electrode (Nano-Z, Neuralynx, MO, USA). Rats were sacrificed under deep anesthesia (100 mg/kg ketamine, 10 mg/kg xylazine IP) by transcardiac perfusion of physiological saline followed by 4% formalin. Brains were extracted and immersed in 4% formalin for 24 hours and then stored in 30% sucrose until ready to be sectioned. Tissue was blocked in the flat skull position using a brain matrix (RWD Life Science Inc., CA, USA). Brains with field potential probes were sectioned frozen in the coronal plane at 50μm thick. Brains with single-unit electrodes were paraffin embedded and sectioned 20 μm thick due to diameter difference in wires (processed by Tissue Technology Shared Resource; CCSG Grant P30CA23100). Brain slices were Nissl stained using thionin to identify the course of the electrode tracks. Sections were processed with a slide scanner at 40x magnification (Zeiss, Oberkochenn, Germany; Leica Biosystems, IL, USA). Positions of electrodes were inferred by matching landmarks in sections to the rat atlas (52) when electrode tips could not be identified (**Supp. Fig 3**).

## Acknowledgments

This work was supported by the following awards: Burroughs Wellcome fund 1015644 (to DR), 1R01MH123650 (to DR), start-up funds from the UCSD Department of Psychiatry (to DR), and a training award T32MH018399 (to MFK). Histopathology was performed in part by Tissue Technology Shared Resource supported by CCSG Grant P30CO23100.

## Supplementary Figures

**Supplemental Figure 1:**
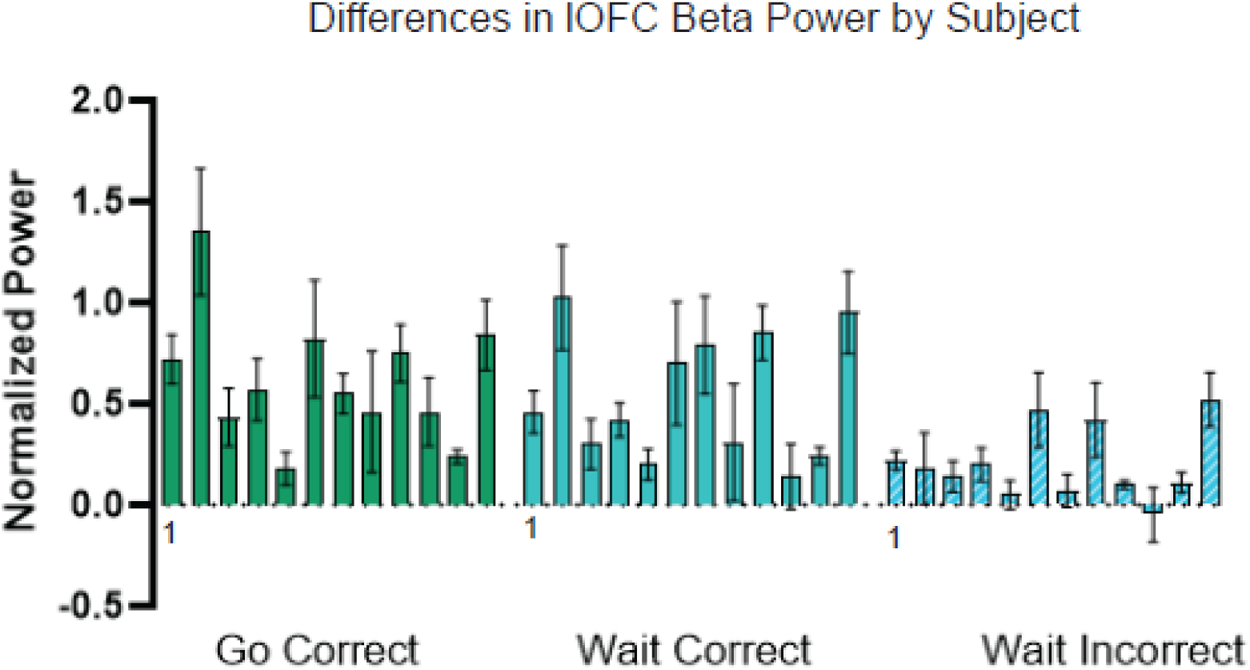
Individual Differences in LOFC Beta Power During Go/Wait Inhibition Task. The average beta power on the LOFC electrode during reward outcome (500-2500ms after response) on go-cue correct (solid green), wait-cue correct (solid blue), and wait-cue incorrect (striped blue) trials for each subject (N=12). Error bars represent SEM.

**Supplemental Figure 2:**
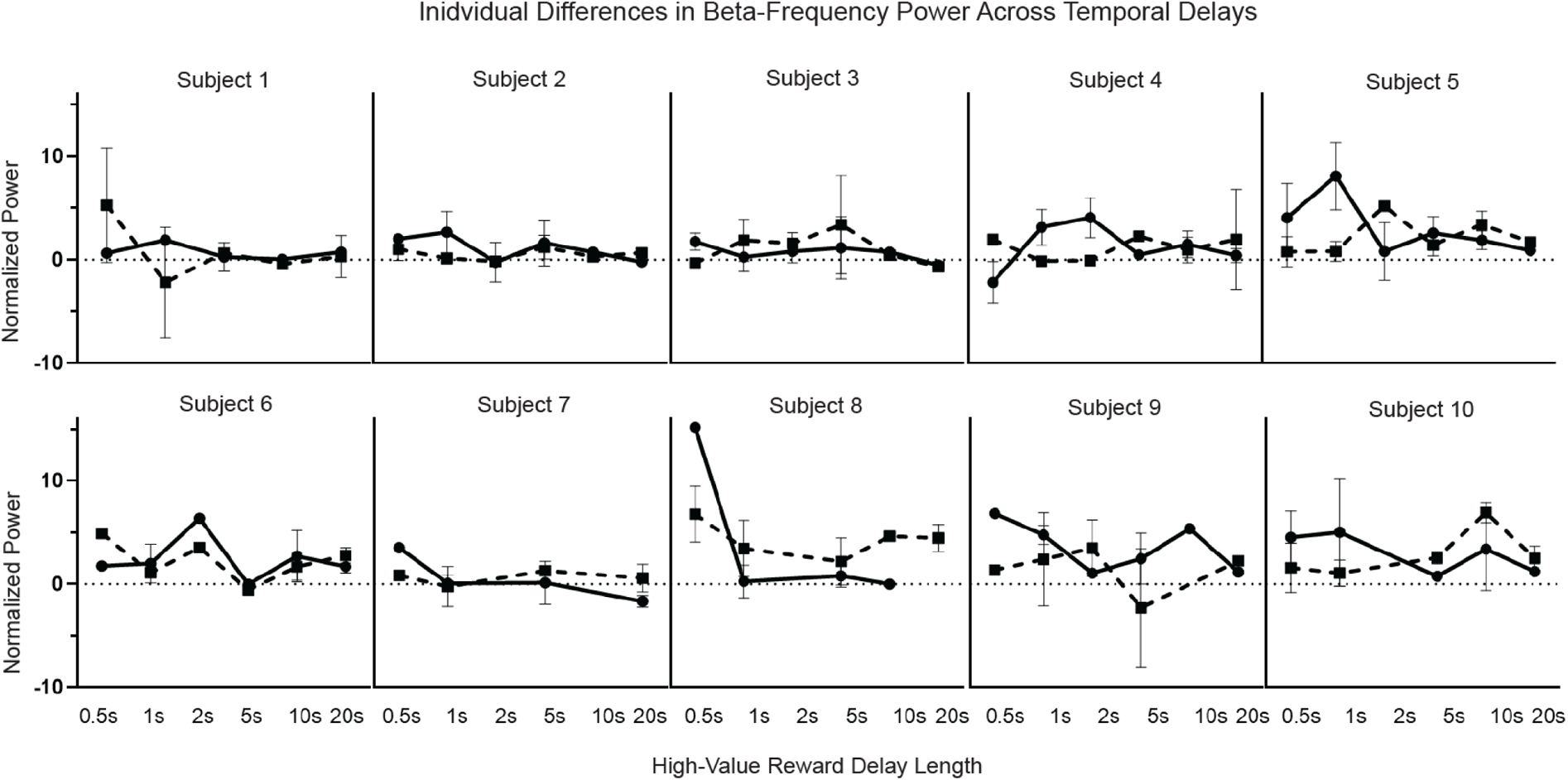
Individual Differences in LOFC Beta Power During Temporal Discounting. The average beta power on the LOFC electrode during reward outcome (0-1000ms after reward) at each temporal delay for each subject (N=10). The two lines represent power on high value choice (solid line) and low value choice (dashed line). Error bars represent SEM.

**Supplemental Figure 3:**
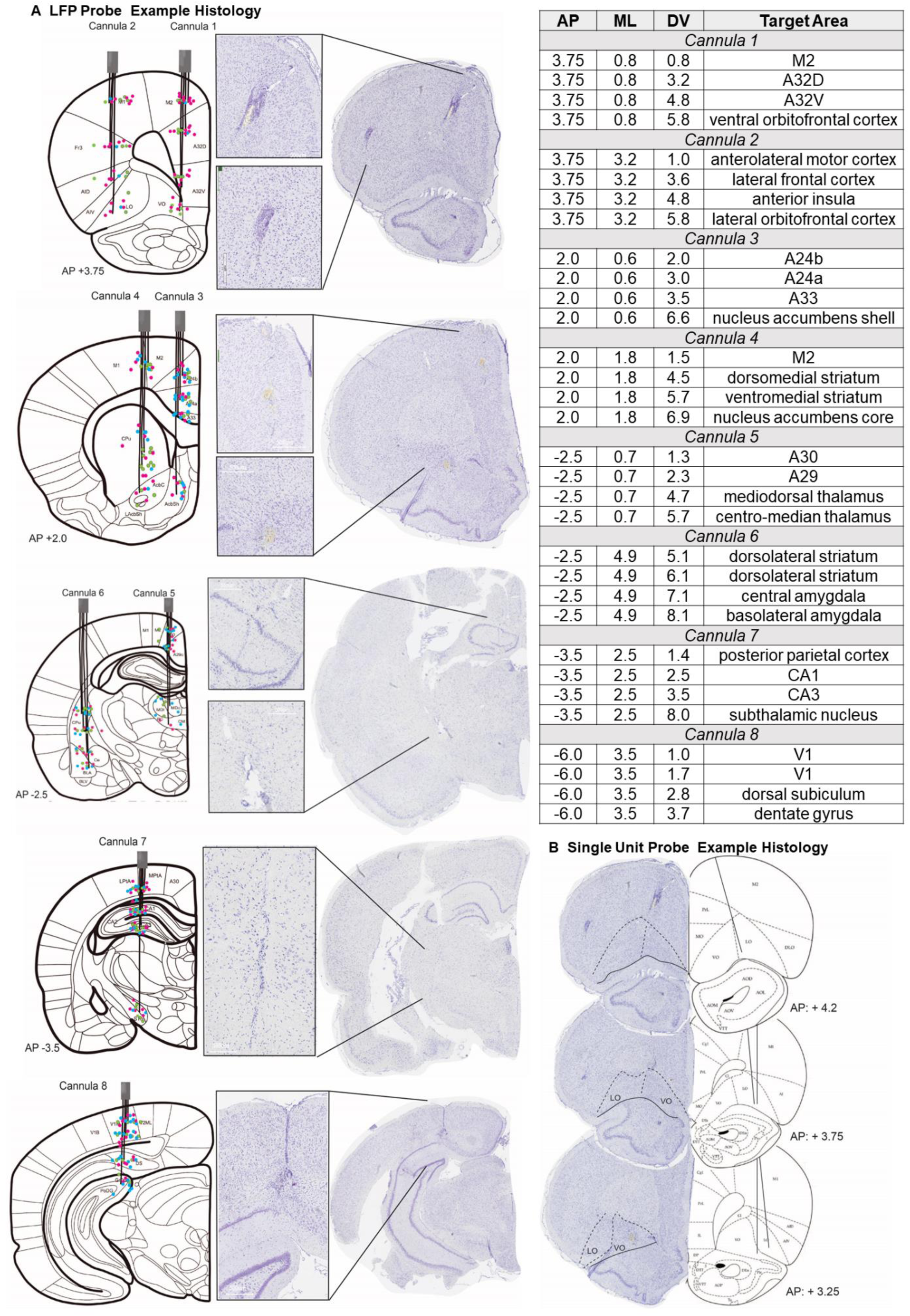
Histological Verification of Recording Sites. (A) LFP implants histology identifying locations of the 32 electrodes (8 cannula). A graphical representation of the placement of each cannula is plotted on a coronal section of a modified rat brain atlas (Paxinos & Watson, 2013) Each cannula contains four wires each targeting a unique DV location. Multiple cannulas may be implanted on the same coronal plane (same AP coordinates). The identified centers of each electrode are shown as dots color coded based on task to (green: go/wait N= 6/11; pink: temporal discounting N= 9/10; blue: PRL N=7/7). An example thionine-stained coronal slice at the corresponding AP location is also shown for each cannula placement with magnification of each track in the brain (white bar provides scale). The table includes the AP, ML, and DV coordinates for all 32 electrodes and their corresponding nomenclature. (B) Single unit implants histology for electrode tracks in OFC. Example coronal sections are shown from +4.2 AP through +3.25 AP relative to bregma. The LO/VO subdivisions are outlined on example thionine-stained slices and the center of the electrode tracks are marked on sections of a modified rat brain atlas (Paxinos & Watson, 2013) (N=5/8 rats).

